# Cell cycle dependent variation in endocytosis drives phenotypic diversity in *M. tuberculosis*

**DOI:** 10.1101/2025.09.09.673684

**Authors:** Neeraja Subhash, Sandhya Krishnan Radhakrishnan, Hitakshi Vijay, Neilay Bhalerao, Sahanawaz Molla, Anton Iyer, Shaon Chakrabarti, Varadharajan Sundaramurthy

## Abstract

Cell-to-cell heterogeneity is a hallmark of biology, yet how host variability shapes intracellular pathogen phenotypes is unclear. Using single cell approaches and redox-sensitive *Mycobacterium tuberculosis* reporters, we reveal that interphase-driven shifts in endocytic capacity create distinct intracellular niches. Bacilli residing in high-endocytic G2 cells adopt more oxidized redox states, whereas those in low-endocytic G1 cells remain reduced, generating phenotypic diversity within a single infected population. Experimental manipulation of host cell cycle stage reprograms endocytic capacity and intrabacterial redox, establishing a causal link between host state and pathogen diversification. Notably, the finding that the cell cycle regulates endocytic capacity constitutes a fundamental cell-biological discovery with broad implications. This variability persists in post-differentiated macrophages, indicating that proliferative history imprints functional diversity onto innate immune cells. Together, these results identify interphase-regulated endocytosis as a host-intrinsic mechanism that shapes *Mycobacterium tuberculosis* phenotypes and suggest new host-directed avenues to influence infection trajectories and persistence.

## Introduction

Cellular heterogeneity is a fundamental feature of all biological systems. Individual cells within a population exhibit substantial variability in their responses despite being in identical environmental conditions. These responses influence processes such as infections,drug susceptibility, stem cell differentiation and cancer progression (Huang, 2009; Altschuler et al., 2010; Dey et al., 2014; Gough et al., 2014; Gingold et al., 2015; Krieger et al., 2015; Komin et al., 2017). Such variability can arise from stochastic transcriptional bursting (Harper et al., 2011; Rodriguez et al., 2019), variation in chromatin states (Carter et al., 2021; Cheow et al., 2016; Linker et al., 2019) and asymmetry during cell division (Q. Chen et al., 2018). A central question is how such heterogeneity arises and whether it acts as a regulatory mechanism or an emergent consequence of cellular complexity.

Host-pathogen interactions are a striking case of heterogeneity, given the interplay of two dynamic systems. During intracellular infection, clonal host cells can display diverse outcomes ranging from variable infectivity, resistance, containment, clearance, or apoptosis - at the same time (Avraham and Hung, 2016; Delince et al., 2016; S. Helaine et al., 2010; McIntrye et al., 1967; Monack et al., 1996). Whether this diversity reflects intrinsic stochasticity, stable cellular states, or dynamic adaptation remains unresolved, yet it likely shapes population-level infection trajectories. Two nonexclusive models have been proposed for variability in host-pathogen encounters: infection-induced heterogeneity versus pre-existing differences that set trajectories from the outset. Single-cell RNA-seq of Salmonella-infected macrophages supports the latter, showing that pre-existing bacterial expression states influence macrophage responses (Avraham, Haseley, et al., 2015). Variation in macrophage polarization creates intracellular niches that allow *Salmonella* either to evade detection and persist or to exploit the host for rapid replication (Saliba et al., 2016). Consistent with this view, our prior work showed that pre-existing heterogeneity in host endocytic capacity drives *M. tuberculosis* (*Mtb*) infectivity and subcellular trafficking (Sachdeva et al., 2020). Likewise, pre-existing differences among primary human vascular endothelial cells (HUVECs) predict susceptibility to *Listeria monocytogenes*, with variability in bacterial adhesion as a principal determinant (Rengarajan et al., 2020). Together, these studies illustrate how host and bacterial heterogeneity intersect to generate distinct subpopulations with divergent behaviors. *Mtb* infection exemplifies this principle, with variability in both pathogen and host contributing to outcomes across cellular and tissue scales (Cadena et al., 2017).

Bacterial phenotypic diversity is advantageous for survival across host environments (S. Helaine et al., 2010; Adams et al., 2011; Sophie Helaine, Cheverton, et al., 2014; Avraham and Hung, 2016; Liu et al., 2016). In *Mtb*, phenotypic diversification is a well-documented survival strategy, with subpopulations adopting distinct physiological states that favor persistence under stress (Parbhoo et al., 2022; Dhar, J. McKinney, et al., 2016). Reporter strains that sense redox, pH, and ATP fluctuations reveal heterogeneous microenvironments within host cells and tissues (Tan, Sukumar, et al., 2013; Bhaskar et al., 2014; Sukumar et al., 2014; Mehta et al., 2015; Tan, Yates, et al., 2017; Akela et al., 2021), and show substantial variability in bacterial states even within the same host context (Bhaskar et al., 2014). While host cellular and tissue environments clearly influence bacterial states and antibiotic recalcitrance (Bhaskar et al., 2014; Sophie Helaine, Conlon, et al., 2024; Liu et al., 2016), the plasticity of *Mtb* phenotypic states and the mechanisms at the host–pathogen interface that drive them remain open questions.

Endocytosis is a fundamental cellular process central to nutrient uptake, signaling, and immune defense, which also varies markedly at the single-cell level. Multiple pathways can operate in the same cell (Mayor et al., 2007; Thottacherry et al., 2019), yet individual cells differ in cargo internalization, vesicle trafficking, and intracellular sorting (Dey et al., 2014). A Genome-wide screen has identified genetic modulators that differentially influence endocytosis, with some exerting pathway-specific effects and others broadly regulating multiple uptake mechanisms (Collinet et al., 2010). Rab5, a key early endosomal regulator, coordinates these processes, and its depletion disrupts both endocytosis and lysosomal function (Zeigerer et al., 2012). While some studies suggest that stochastic fluctuations and cell–cell communication shape these differences (Altschuler et al., 2010; Snijder et al., 2009), the origins and drivers of endocytic heterogeneity remain unclear. Given that *Mtb* primarily enters host cells via phagocytosis and our previous work linking endocytic heterogeneity to *Mtb* infectivity and subcellular trafficking (Sachdeva et al., 2020), understanding how subpopulation-level differences in endocytosis affect infection dynamics is particularly relevant. Single-cell RNA-seq and live-cell imaging have revealed substantial host and bacterial heterogeneity during infection (Mills et al., 2017; Delince et al., 2016; Mottet et al., 2021), while *Mtb* stress-reporters (Akela et al., 2021; Bhaskar et al., 2014; Mehta et al., 2015; Sukumar et al., 2014; Tan, Sukumar, et al., 2013; Tan, Yates, et al., 2017) expose coexisting bacterial subpopulations in distinct host niches. Yet, we lack a temporal, causal framework linking specific host cell states to the emergence and dynamics of these bacterial phenotypes.

We integrate live-cell imaging, scRNA-seq, and redox-sensitive *Mtb* reporter to determine how host cell heterogeneity drives bacterial phenotypes. We show that cell-cycle progression during interphase is a major source of single-cell variation in endocytic capacity, and that these differences drive phenotypic diversity in intracellular *Mtb*. Cells with distinct endocytic capacity harbor *Mtb* in different redox states, and experimental manipulation of the cell cycle reprograms both host endocytic profiles and the distribution of bacterial phenotypes. Temporal shifts in endocytic capacity during interphase are mirrored by corresponding changes in intracellular *Mtb* states, establishing a dynamic, causal link from host cell state to pathogen heterogeneity. Thus, interphase progression regulates endocytic heterogeneity and actively shapes the bacterial phenotypic distribution.

## Methods

### 1 Cell Culture

RAW 264.7, THP-1 monocytes, and J774 cell lines were cultured in RPMI medium (Gibco, 31800022). HeLa, HEK293FT, immortalised BMDMs, and Vero E6 cells were maintained in DMEM (Gibco, 12800017). Both media were supplemented with 10% fetal bovine serum (heat-inactivated) (FBS; Gibco, 16000044) and 1× Antibiotic-Antimycotic (100×; Gibco, 15240062). THP-1 monocytes were differentiated into macrophages using 20 ng/mL phorbol 12-myristate 13-acetate (PMA; Sigma-Aldrich, P8139) as described in (Baxter et al., 2020). HeLa BAC GFP-Rab5 cells were cultured in complete DMEM supplemented with 500 *µ*M Geneticin (G418; Invitrogen, 10131035). L929 cells were maintained in complete DMEM.cell-conditioned medium was collected from confluent L929 cultures.All cells were maintained at 37*^◦^*C in a humidified atmosphere containing 5% CO_2_ and 95% relative humidity.

#### Isolation of Bone Marrow Derived Macrophages (BMDMs) for infection and functional assays

Monocytes were isolated from the bone marrow of C57BL/6 mice and differentiated into macrophages following a standard protocol (Toda et al., 2021; Nagabhushanam et al., 2003). Briefly, bone marrow was flushed from the tibia and femur using a syringe filled with complete DMEM, and cells were plated in DMEM supplemented with 10% fetal bovine serum (FBS) and 20% L929-conditioned medium. After 7 days of differentiation, cells were washed with PBS and seeded into 12-well plates. Subsequently, these macrophages were used for infection experiments and functional assays, including dextran uptake and cell cycle analysis using DRAQ-5.

#### Endocytic Uptake Assay

Endocytic uptake assays were performed using fluorescently labeled dextran (Life Technologies, D-1817 and D-22914) or transferrin (Life Technologies, T-2336 and T-23365). Cells were pulsed with dextran (1 mg/mL) or transferrin (5 µg/mL) for 10 minutes in incomplete RPMI or DMEM as described in (Sachdeva et al., 2020). Following the pulse, cells were washed with incomplete RPMI and processed for downstream assays.

#### Sorting cells into subpopulations based on endocytic capacity

Cells from the parental endocytic population were sorted into low (10–15%) and high (10–15%) endocytic subpopulations based on the fluorescent intensity of the cargo or GFP-Rab5 intensity in a BD FACS Aria Fusion or Aria III sorter. After sorting, cells were analyzed to confirm the accuracy of the fluoresence intensity distribution of the sorted subpopulations. The sorted subpopulations were maintained in RPMI or DMEM media supplemented with 10% fetal bovine serum (FBS) at 37°C in a humidified atmosphere containing 5% CO_2_ and 95% relative humidity.

#### Live cell imaging and image analysis

Live-cell imaging of the ‘low’ and ‘high’ endocytic subpopulations of HeLa BAC GFP-Rab5 cells was performed using a NIKON Ti2 Eclipse epifluoresence microscope equipped with an on-stage incubator (OKO LAB) to maintain physiological conditions (37°C, 5%CO_2_, and 95% relative humidity) for 72 hours, with imaging intervals of 30 minutes. We built a customised image analysis pipeline. First, image pre-processing, including background subtraction, was performed using Fiji (Schindelin et al., 2012). Cells were segmented using the GFP fluorescence using CellPose (version 2.3.2) (Stringer et al., 2021; Pachitariu et al., 2022), with a custom segmentation module trained on the Cyto2 model using over 2000 manually curated masks. All segmentation masks were manually verified, and erroneous masks corrected as needed. The resulting cell masks were imported into Ilastik (version 1.4.0) (Sommer et al., 2011) for lineage tracking. All lineage tracks were manually verified. An example of segmentation masks and lineage tracking is provided in Figure S3. Intensity profiles and lineage data were exported from ilastik as CSV files and analyzed in R, using the ggplot2 and dplyr libraries for data visualization and analysis.

#### Cell synchronization

HeLa BAC GFP-Rab5 cells were seeded in 12-well plates at 60% confluency and treated the following day with 2.5mM thymidine for 18 hours. After treatment, cells were washed to remove thymidine (release) and cultured in complete DMEM for 8 hours. At 0 or 8 hours post-release, cells were pulsed with transferrin-AF647 for 10 minutes, washed three times with incomplete media, trypsinised, and fixed in 80% ethanol. Cell cycle analysis was performed using FlowJo v10.8 following a modified protocol from (Kim et al., 2015). Briefly, ethanol fixed cells were then centrifuged at 3500 RPM (1150*g) for 5 minutes, and resuspended in PBS containing 0.1 % Triton X-100 and Hoechst (5 µg/mL). Cells were centrifuged after 10 minute incubation and resuspended in FACS buffer (1% FBS in 1X PBS) and analysed using BD FACS Fortessa X20 SORP. Cell cycle analysis of RAW 264.7 cells was similar to the above, with the following modifications: thymidine treatment lasted 12 hours, cells were fixed with 4% PFA for 4 hours after release from the block, and then removed from the culture dish by scraping.

#### Immunostaining, imaging and analysis

RAW 264.7 macrophages and HeLa cells were fixed with 4% paraformaldehyde and permeabilized in SAP buffer (0.2% saponin (Sigma-Aldrich, S4521), 0.2%gelatin (HML, TC041) in PBS) for 10 minutes at room temperature. Cells were then incubated with primary antibodies against EEA1 (1:100; CST #3288 or BD 610457) and RAB5 (1:100; BD 610282) for 2 hours at room temperature. After washing with SG-PBS (0.02% saponin, 0.2%gelatin in PBS), cells were incubated with appropriate secondary antibodies prepared in SG-PBS (1:500) for 1 hour at room temperature, following which cells were washed with SG-PBS and stained with DAPI (1 µg/mL; Invitrogen, D1306) for nuclear visualization. Imaging was performed using either a Nikon Ti2 or Opera Phenix microscope. Image analysis was performed using CellProfiler(Kamentsky et al., 2011) or Harmony software, using pipelines adapted from previous publication (Sachdeva et al., 2020). Briefly, cells were segmented to identify nuclei, cell boundaries, and endosomal compartments. Segmented objects such as nuclei and endosomes were assigned to individual cells, and multiple features—including number, size, and fluorescence intensity—were extracted at single-cell level. Data analysis and visualization were performed in RStudio (Version 2024.04.1+748) using the ggplot2 and dplyr packages.

#### *M.tuberculosis* Mrx1-roGFP2 H37Rv Infection

*Mycobacterium tuberculosis* cultures were grown in Middlebrook 7H9 broth supplemented with Middlebrook OADC Enrichment (BD Biosciences, 211886), 5% glycerol, and 0.05% Tween-80 (Sigma-Aldrich, P4780). For all experiments, bacterial cultures were used in mid-log phase (OD_600_ = 0.6–0.8). Hygromycin B (Invitrogen, 10687010) was added at a final concentration of 50 µg/mL for the maintenance of the *M. tuberculosis* Mrx1-roGFP2 H37Rv strain.

The *M. tuberculosis* Mrx1-roGFP2 H37Rv reporter strain was used to assess intracellular redox potential in ‘low’ and ‘high’ endocytic subpopulations. Macrophages (RAW, THP-1 monocyte derived, or bone marrow derived) cells were seeded in 12-well plates at a density of 4 × 10^5^ cells per well and infected with a multiplicity of infection (MOI) of 10 for 2 hours. Cells were washed to remove extracellular *Mtb* and pulsed with either Transferrin-AF647 or Dextran-TMR for 10 minutes, followed by fixation with N-ethyl maleimide (NEM) for 10 minutes and 4% paraformaldehyde (PFA) for 25 minutes.

Each experiment included calibration controls to measure the response of the *Mtb* reporter strain to standard oxidative (cumene hydroperoxide, CHP; 5 mM) and reductive (dithiothreitol, DTT; 40 mM) treatments. Samples were analyzed by flow cytometry on a BD LSR Fortessa. Emission from *Mtb*Mrx1-roGFP2 was collected at 510 nm following two different excitations at 405 nm and 488 nm, respectively. For each experiment, data were acquired from 10,000 infected cells and analyzed using FlowJo v10.8. For redox measurements of intracellular *Mtb* in endocytic subpopulations, cells were stratified into the lower and upper 30% of the transferrin or dextran fluorescence intensity distribution, representing ‘low’ and ‘high’ endocytic capacity groups, respectively.

Intracellular redox potential was calculated as previously described in (Bhaskar et al., 2014). Briefly, the emission ratio at the two excitation wavelengths provides a quantitative measure of the redox stress experienced by intracellular *Mtb*. The degree of oxidation was determined by normalizing the observed fluorescence ratio to values obtained from oxidized (CHP) and reduced (DTT) controls using the following equation:

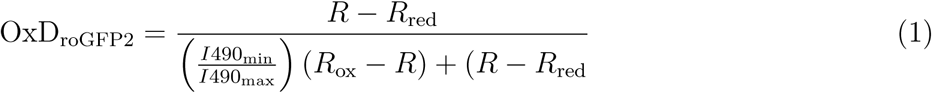

where R is the observed ratio, R_red_ and R_ox_ are the ratios of completely reduced (DTT treatment) and oxidized (CHP treatment) roGFP2, respectively. I 490_min_and I 490_max_ are the fluorescence intensities measured with excitation at 490 nm for fully oxidized and fully reduced roGFP2, respectively. The intracellular redox potential E_roGFP2_ was calculated using the Nernst equation:

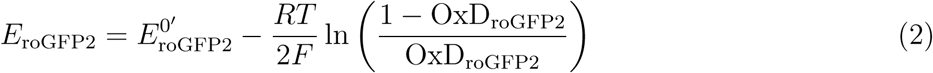

Based on the midpoint redox potential range of roGFP2 (–270 to –280 mV), gates were defined to determine the proportion of basal, oxidized, and reduced E_MSH_ *Mtb*subpopulations. For each experimental condition, the percentage of cells within each E_MSH_ subpopulation was calculated and visualized using Microsoft Excel.

To assess bacterial redox states across different cell cycle stages, RAW 264.7 cells were pre-treated with thymidine (2 mM) for 12 hours. Infections were carried out at 0 and 4 hours post-release using *Mtb* Mrx1-roGFP2 H37Rv at an MOI of 20 for 30 minutes. The high MOI in these experiments was necessitated by shorter infection times due to active cycling of RAW cells. Following infection, cells were fixed with NEM for 10 minutes and then with 4% PFA for 25 minutes. For cell cycle analysis, the standard Hoechst could not be used due to *Mtb* fluorescence at 405 nm, instead DRAQ-5 (Cell Signaling Technology, #4084) was used at a 1:500 dilution for 10 minutes at room temperature prior to FACS analysis. To assess bacterial redox states between cells at different cell cycle stages, measurements were compared between untreated and thymidine-treated cells.

#### Single cell RNA sequencing

To check the differential gene expression at single cell level between endocytic subpopulations, RAW 264.7 cells were pulsed with dextran-TMR (1 mg/ml) for 10 mins and sorted into low 10-15%and high 10-15% endocytic subpopulations. In parallel, cells from the same passage were treated with 2 mM thymidine for 12 hrs, the block released and the cells were grown in complete RPMI for 3hrs 30 min. For the single cell sequencing, the sorted low and high dextran subpopulations, along with parental and thymidine treated cells (after 3hrs 30 min of release) were taken as different samples. All the samples were tested for cell viability prior to single cell sequencing. Sequencing was performed using Chromium Next GEM Single Cell 3’LT Kit v3.1 (PN-1000325)(User Guide CG000399). Briefly, cells were resuspended in the RT master mix and loaded into a Next GEM Chip G, along with gel beads and partitioning oil, to generate Gel Bead-In-Emulsions (GEMs) using the Chromium Controller. Within each GEM, cells were lysed and polyadenylated RNA was reverse transcribed, incorporating a template switch oligo and barcoded oligo-dT primers that included an Illumina Read 1 (R1) primer sequence, a 10x cell barcode, and a Unique Molecular Identifier (UMI). Following reverse transcription, GEMs were broken and the pooled, barcoded cDNA was purified using Silane DynaBeads, then amplified by PCR. Size selection of appropriately sized cDNA fragments was performed using SPRIselect beads. During library construction, enzymatic fragmentation, end repair, and A-tailing were followed by adapter ligation and PCR amplification to add the Illumina Read 2 (R2) primer site, P5/P7 flow cell adapters, and a sample index. The final libraries were assessed for quality and size prior to sequencing on an Illumina platform. The data were analysed following current best practices for scRNA-seq analysis ((Heumos et al., 2023) as detailed below.

#### scRNA seq Data Analysis

##### Single-cell RNA-seq Preprocessing

Raw FASTQ files were processed using Cell Ranger v7.2.0 (10x Genomics) on a Linux system. The software package (`cellranger-7.2.0.tar.gz`) and the mouse reference genome (`refdata-gex-mm10-2020-A.tar.gz`) were extracted using the `tar` command. The reference transcriptome used was *Mus musculus* mm10 (2020-A release). The `cellranger count` pipeline (v7.2.0, 10x Genomics) was run individually for each sample using appropriate parameters to process demultiplexed FASTQ files against the `mm10-2020-A` reference genome. This step performed alignment, barcode and UMI processing, and gene expression quantification, generating gene–barcode matrices and associated quality control metrics for downstream analysis.

###### Data Import and Construction of AnnData objects

To enable downstream single-cell RNA-seq analysis in Python, the Cell Ranger output matrices (matrix.mtx.gz, features.tsv.gz, and barcodes.tsv.gz) were parsed and converted into a gene-by-cell expression matrix. This matrix was saved as a CSV file and subsequently used to construct an AnnData (.h5ad) object, incorporating both gene and cell metadata, facilitating further analysis using the Scanpy framework (Wolf et al., 2018).

###### Quality Control in Python

Quality control (QC) metrics were calculated to assess cell quality across all conditions by identifying mitochondrial genes (prefix `mt-`) in gene annotations. Using Scanpy’s `calculate qc metrics` function, we computed standard QC metrics, including the proportion of mitochondrial reads, the number of highly expressed genes, and logarithmically transformed features. This ensured the proper identification of cells with potential mitochondrial contamination.

In addition to cell-level filtering, genes were filtered based on their expression levels. Specifically, only genes expressed in at least 20 cells were retained for downstream analysis. This step reduces technical noise and focuses the analysis on biologically relevant signals.

Violin plots were generated for key QC metrics: number of genes detected (`n genes by counts`), total counts (`total counts`), and percentage of mitochondrial genes (`pct counts mt`) for both high and low conditions. These visualizations helped identify low-quality cells exhibiting characteristics such as low gene detection, high mitochondrial content, or outlier total counts.

Based on these plots, cells with excessive mitochondrial contamination (greater than 10%) and outliers in gene count or total counts were filtered out to improve overall data quality.

###### Normalization and Log Transformation

To enable comparison between conditions, a new column labeled `Condition` was added to the `obs` metadata in all datasets. The data were then normalized using the `normalize total` function, scaling counts per cell to 10^4^ to account for varying sequencing depths. A log transformation (`log1p`) was applied to stabilize variance and reduce the influence of highly expressed genes. Finally, the processed datasets were saved as `.h5ad` files for future analysis.

###### Data concatenation and Pseudobulk generation

After quality control, the datasets to be analysed for differential gene expression were merged into a single `AnnData` object using `anndata.concat()` along the cell axis, with a new `Condition` column added to indicate the condition (“high” or “low”). The combined dataset was saved as `combined quality control.h5ad` for downstream analysis. For pseudobulk analysis, the count matrix was extracted from the combined `AnnData` object as a `pandas DataFrame` with cell barcodes made unique and condition labels retained. To generate the pseudobulk data, cells from each condition were randomly grouped into three pseudoreplicates using a fixed random seed for reproducibility. The resulting annotated matrix, containing both `Condition` and `Replicate` columns, was saved along with the corresponding cell-level metadata and gene-level metadata for use in downstream analyses.

###### Data Aggregation and Dimensionality Reduction

The data was then aggregated by summing expression values across all cells within each replicate for both conditions, creating replicate-level profiles. These profiles were saved as new AnnData objects and later combined into one dataset.

###### Differential Gene Expression Analysis

To identify differentially expressed genes (DEGs) between conditions, the pseudobulked AnnData object was analyzed using the edgeR package in R via the rpy2 interface. Genes with a false discovery rate (FDR) less than 0.05 were considered significant. All intermediate R objects and results, including the fitted model, DEG list, design matrix, and statistical results, were saved as RDS files to ensure reproducibility and facilitate future analysis.

#### Single cell colony sorting and expansion

Single-cell sorting was performed using a Beckman CytoFLEX SRT three-laser system. HeLa GFP-RAB5 cells were cultured to confluency, and GFP-positive cells were gated using wild-type HeLa cells as a negative control. Individual GFP-positive HeLa cells were sorted into 384-well optical bottom plates containing 80 µL of DMEM supplemented with 10% FBS and 1× Antibiotic-Antimycotic (100×). Following sorting, cells were cultured for 3 days, after which the media was replenished. After one week, wells with visible cell colonies were trypsinized and expanded into 48-well plates and subsequently into 25 cm² flasks. Once sufficient expansion was achieved, stocks for each clonal population were generated from cells cultured in T25 flasks. These clonal stocks were then used for downstream experiments.

#### Determination of Asymmetry

To determine the coefficient of asymmetric binomial partitioning of GFP-Rab5 in the daughter cells from the time-lapse data, cells were classified into mothers and daughters. The analysis was based on the relationship of the variance in intensity of daughter cells predicted from the statistics of the binomial distribution process (Peruzzi et al., 2021).

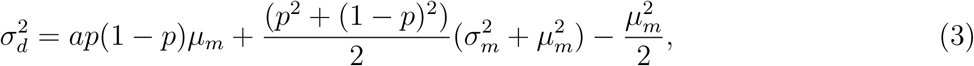

where *σ_d_* is the standard deviation of GFP RAB5 intensities in the daughter cells just after birth, *a* is a rescaling variable to rescale the GFP RAB5 intensities by 10*^−^*^6^, *p* is the probability of partitioning, *µ_m_* and *σ_m_* is mean and standard deviation of the scaled GFP RAB5 intensity in the mother cells just before division. Cells tracked for 5 hours or more were selected for analysis. For these cells, mother and daughter identities were assigned. The mean and standard deviation of total GFP-Rab5 intensity (TI) were calculated separately for mothers and daughters. For mothers, the TI value was taken 30 minutes prior to cell division; for daughters, the TI value was recorded at the birth timepoint (i.e., immediately after division). All calculations were carried out independently for each of the three strains (High, Parent, Low).

To estimate the asymmetric binomial partitioning coefficient *p*, a bootstrapping approach was employed. TI data were sampled with replacement 1000 times. In each iteration, a mean and standard deviation of TI were computed for mother and daughter cells, which were then used to calculate a value of *p* using the partitioning equation (Eq.3). Repeating this over 1000 iterations produced a distribution of *p* values.

#### Comparison of Inter-mitotic Times (IMT) of cells across strains and generations

Cells in different strains were segregated into generations based on frequency of the divisions. Starting cells were categorized as generation ‘zero’, their progeny were subsequently termed as ‘first’, ‘second’ and so on. For the generation zero cells, the time to their division was extracted and used as a proxy for IMT. For the rest of the generations, the time interval between birth and the division event was used to calculate the IMT. Then, the IMT of cells belonging to different strains was compared for all the generations. A non parametric Wilcoxon Rank Sum Test was performed to compare the distributions as the data was non-Gaussian with long tails.

#### Linear regression for calculating slopes of time-series data

Time-lapse imaging of HeLa BAC cells provided repeated measurements of GFP–Rab5 intensity in the same cells over time under different experimental conditions. Since total GFP intensity increased monotonically, we applied a linear model (*lm* function in R) to estimate the slopes. For untreated and post–thymidine block cells, track lengths were trimmed to match the thymidine block. Although the block itself was 18 h, we used a 20-h analysis window to allow 2 h for cell recovery. Tracks shorter than 3.5 h were excluded. Data analysis and plotting were carried out in KNIME and R.

#### Variance decomposition analysis

The contribution of a random variable *X* to the total variance of a random variable *Y* can be computed as follows:

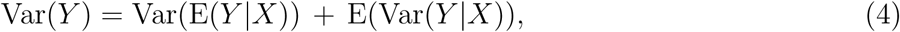

where the first term on the right hand side denotes the variance explained by *X*, while the second term corresponds to the remnant variation in *Y* that cannot be explained by *X*. In our context, *Y* is either GFP-Rab5 or Transferrin intensity, while the covariate *X* is Hoechst intensity. Channel values from flow cytometry data was used for these calculations.

## 2 Results

### Heterogeneity in endocytic capacity promotes *Mtb* phenotypic diversity

Our previous work showed that macrophage subpopulations with higher endocytic capacity (‘high’ cells) are more permissive to *Mtb* infection than those with lower capacity (‘low’ cells). Moreover, *Mtb* trafficking differs between these groups, with bacteria in ‘high’ cells more frequently delivered to lysosomes (Sachdeva et al., 2020), suggesting that they may encounter greater lysosome-associated stress. Such variation in endocytic capacity could therefore generate phenotypic diversity in *Mtb*. To test this, we infected RAW macrophages with *Mtb* expressing Mrx1-roGFP2, a redox-sensitive reporter that enables fluorescence-based measurement of oxidative stress associated with phago-lysosomal compartments (Bhaskar et al., 2014). Based on the ratio of bacterial fluorescence intensities at 405 nm and 488 nm excitation, infected cells were classified as harboring E_MSH_-‘oxidized,’ ‘reduced,’ or ‘basal’ *Mtb* phenotypes (Bhaskar et al., 2014). Following infection, cells were pulsed with fluorescently tagged dextran or transferrin - endocytic cargoes with distinct uptake routes and fates (Sachdeva et al., 2020). Fluorescence intensities of these cargoes were then used to define ‘high’ or ‘low’ endocytic capacity subpopulations, and the infectivity and redox state of intracellular *Mtb* were assessed in these groups (schematic, Figure 1A). Consistent with our previous findings (Sachdeva et al., 2020), the ‘high’ endocytic capacity subpopulations showed higher infectivity (Figure1B). Importantly, these subpopulations also harbored a significantly greater proportion of E_MSH_-oxidized *Mtb* compared to ‘low’ endocytic capacity subpopulations. Conversely, the proportion of cells harboring E_MSH_-reduced *Mtb* was significantly lower in the ‘high’ group (Figure 1B; FigureS1A). Similar results were observed in infections of THP-1 monocyte-derived macrophages (Figure1C) and primary mouse bone marrow–derived macrophages (Figure1D). These findings indicate that intracellular *Mtb* in cells with higher endocytic capacity experience greater oxidative stress than in those with lower capacity. Across independent experiments and infection systems, high endocytic capacity was consistently associated with a decrease in the E_MSH_-reduced subpopulation and a corresponding increase in the E_MSH_-oxidised subpopulation, indicating a robust redistribution from reduced to oxidised states. The E_MSH_-basal fraction showed only minor changes, suggesting that high endocytic capacity primarily drives exit from the reduced state and promotes oxidised phenotypes without substantially altering the transitional pool. Together, these results demonstrate that host endocytic capacity shapes phenotypic heterogeneity and stress adaptation in intracellular *Mtb*.

**Figure 1:**
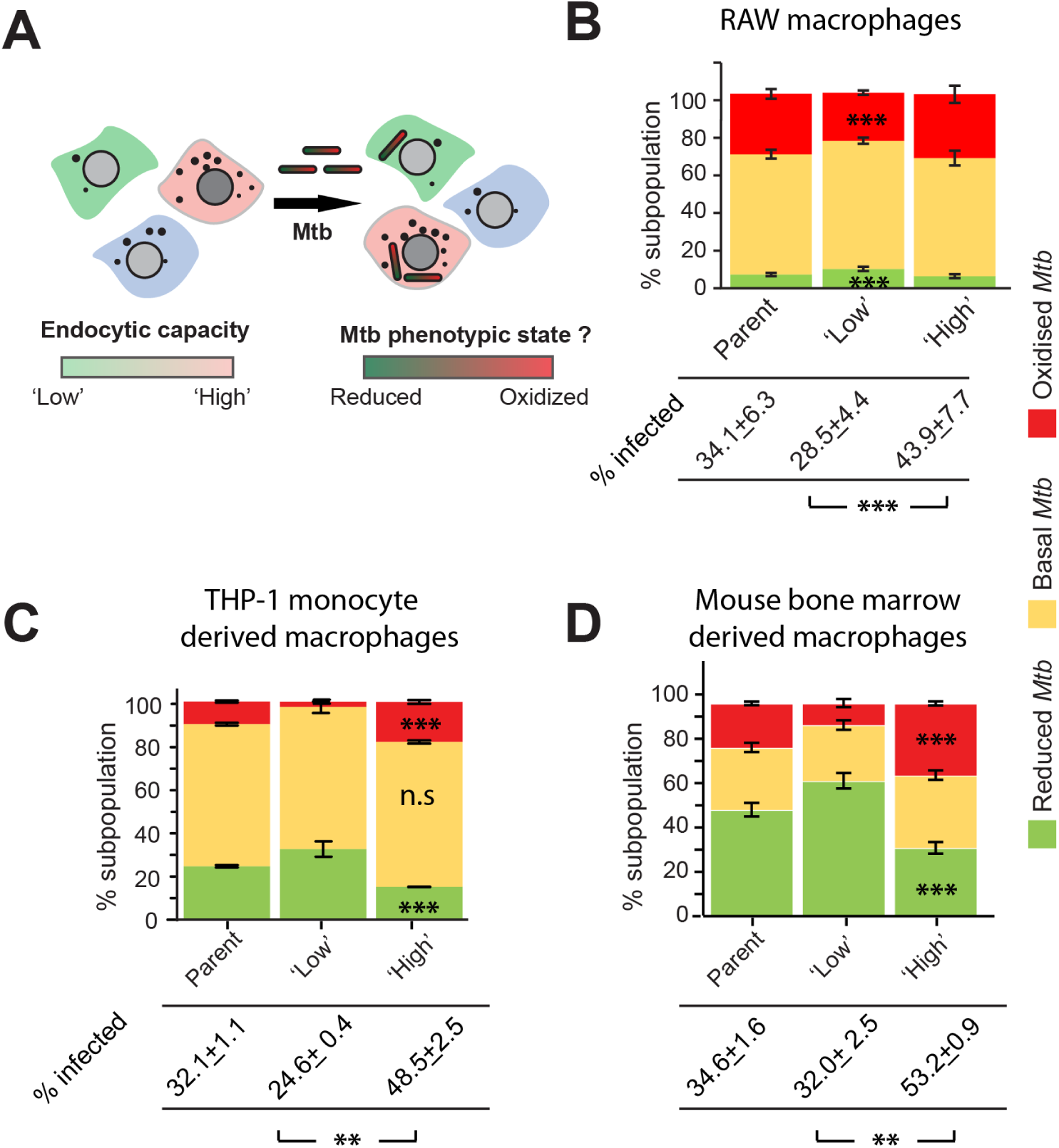
Heterogeneity in endocytic capacity promotes *Mtb* phenotypic diversity. **A**) Schematic of experiment to assess *Mtb* redox states in cells of different endocytic capacities. RAW 264.7 macrophages were infected with *Mtb* Mrx1-roGFP2 H37Rv and pulsed with dextran-TMR for 10 minutes. Cells were then fixed and analyzed by flow cytometry. Based on cargo fluorescence intensity, infected cells were categorized into ‘high’ or ‘low’ endocytic capacity subpopulations, and the redox status of intracellular *Mtb* was assessed within each group. A total of 10,000 infected cells were analyzed, with the lower and upper 30% of the dextran-TMR distribution used to define the ‘low’ and ‘high’ endocytic subpopulations, respectively. The proportions of bacterial subpopulations within each host subpopulation were calculated as described in the Methods. **B**) Infected cell subpopulations defined based on dextran uptake harbour *Mtb* of different redox states denoted as E_MSH_-reduced, basal and oxidised. Percentage of infected cells across these host subpopulations are indicated. **C,D**) E_MSH_-reduced, basal and oxidised *Mtb* in THP-1 monocyte derived macrophages (C) or mouse bone marrow derived macrophage (D) subpopulations defined by dextran uptake. Data the representative of at least three independent biological experiments. Error bars represent standard deviation from the mean of technical replicates. *denote *p*-values from Student’s *t*-test (paired):\*\**p <* 0.01, \*\*\**p <* 0.001; ns, not significant.

### Characterization of variance in the endocytic capacity across cell types

Our findings raised the question of why endocytic capacity itself varies across cells. To address this, we next investigated the sources of this variation as a cell-intrinsic property, independent of infection. Rab5, a key regulator of endocytosis (Zeigerer et al., 2012), is essential for early endosome formation. Overexpression of Rab5 accelerates both fluid-phase and receptor-mediated endocytosis, establishing it as a rate-limiting factor in this pathway (Bucci et al., 1992). Based on this, we hypothesized that total Rab5 levels determine the endocytic capacity of a cell. To test this, we sorted RAW macrophages into ‘high’ and ‘low’ subpopulations based on dextran uptake (Figure 2A) and immunostained for Rab5 and its effector, Early Endosomal Antigen 1 (EEA1). Image analysis revealed that the number and intensity of Rab5- and EEA1-positive endosomes co-varied with dextran uptake in the ‘high’ and ‘low’ subpopulations (Figure 2B). Thus, early endosomal content, together with cargo uptake, defines the endocytic capacity of distinct subpopulations. We then asked whether this capacity is stable or dynamic over time. Sorted RAW macrophage subpopulations were cultured for three days, and their endocytic capacity was re-analyzed. Notably, the initially distinct ‘high’ and ‘low’ populations converged over time (day 0 to day 3), becoming indistinguishable from the parental distribution (Figure 2C). Similar results were obtained in Raw cells infected with *Mtb* (Figure 6D i, iii). These findings demonstrate that endocytic capacity is not fixed but a dynamic property of individual cells.

**Figure 2:**
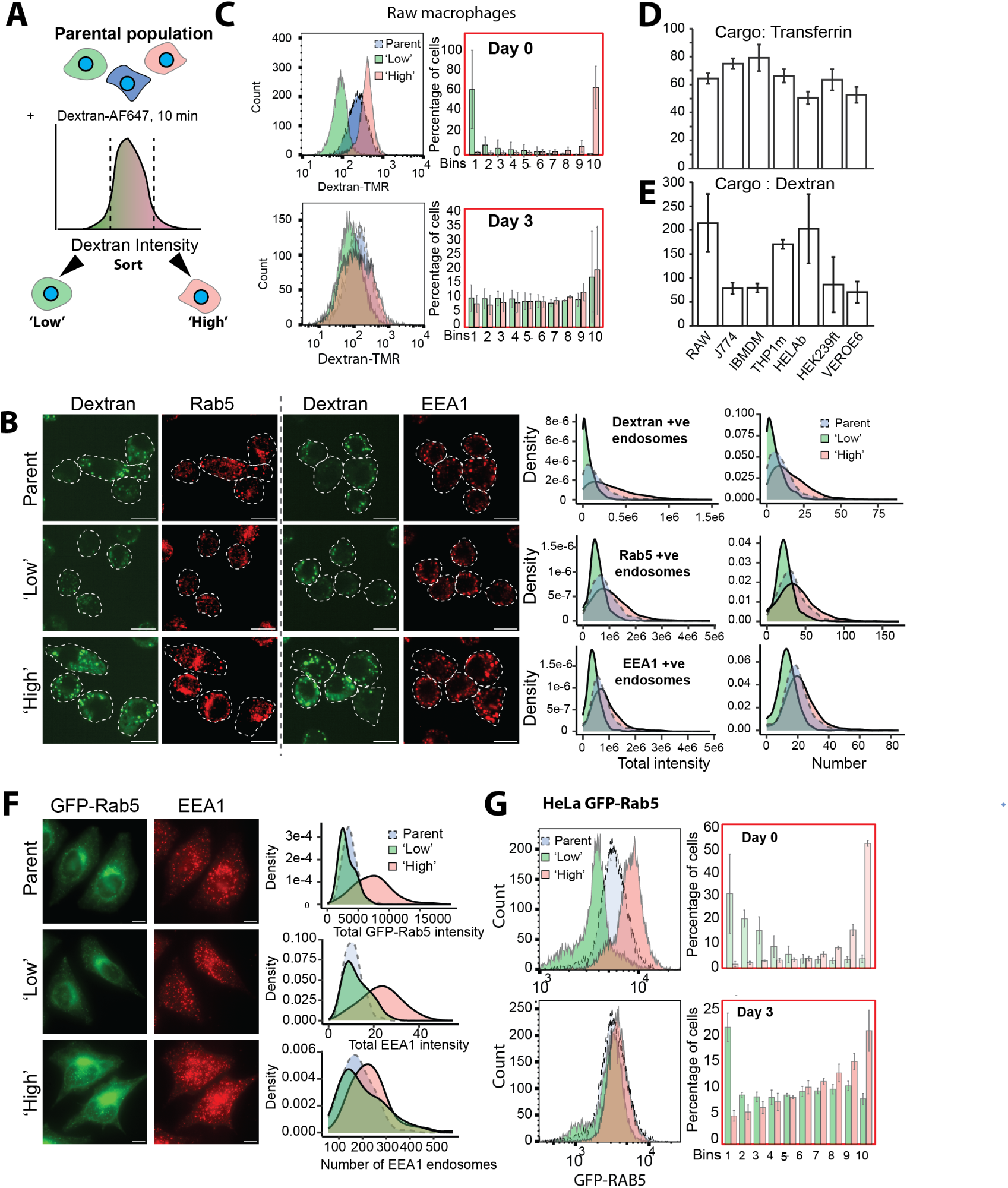
Characterization of variance in the endocytic capacity across cell types. **A**) RAW 264.7 macrophages were pulsed with dextran conjugated to fluorophore for 10 minutes to assess endocytic activity. Cells were then sorted based on fluorescence intensity from the two ends of the distribution : the lowest and highest 10–15 % were designated as ‘Low’ and ‘High’ endocytic capacity subpopulations, respectively. **B**) EEA1 and Rab5 immunostaining in Dextran-TMR sorted Parent, ‘Low’ and ‘High’ subpopulation of RAW 264.7 macrophages, fixed immediately after sorting. Density plots from 800-2500 cells shows the distribution of the number of puncta and total intensity of Dextran, EEA1 and Rab5 endosomes between the subpopulations. **C**) Distribution of dextran uptake from 10,000 cells in parental and sorted RAW 264.7 macrophage subpopulations. Fluorescence intensity profiles of dextran-AF647 in the parental population, as well as in the ‘Low’ and ‘High’ endocytic capacity subpopulations, were assessed either immediately after sorting *(i, Day 0)* or after three days in culture post-sorting *(ii, Day 3)*. Bar graphs represent binning analysis of the percentage of cells from each group falling into defined fluorescence bins, based on the dextran distribution of the corresponding parental population at each time point. Error bars indicate standard deviation from the mean of three independent biological replicates. **D,E**) Dextran or transferrin uptake assay was performed independently in the different cell lines indicated. The coefficient of variation (CV) for dextran-TMR and transferrin-488 was calculated from the fluorescence intensity distributions after background subtraction. Background subtraction was performed by subtracting the mean fluorescence intensity of the unstained control from each event’s scale value. Error bars indicate standard deviation from mean between three biological replicates. **F**) HeLa GFP-Rab5 cells are sorted into ‘low’ and ‘high’ subpopulations based on GFP-Rab5 intensity, fixed and immunostained for EEA1. Representative images show GFP-Rab5 and EEA1 endosomes from parental, ‘low’ and ‘high’ endocytic subpopulations. Density plots from 60-90 cells show the distribution of the number and intensities of the endosomes between the subpopulations. **G**) Distribution of GFP-Rab5 intensity from 10,000 cells ‘low’ and ‘high’ sorted HeLa GFP-Rab5 subpopulations either immediately (i, day 0) or three days (ii, day 3) post-sorting. Black dots denote the parental distribution. Bar graphs represent binning analysis of the percentage of cells from each group falling into defined fluorescence bins, based on the GFP-Rab5 distribution of the corresponding parental population at each time point. Error bars indicate standard deviation from the mean of three independent biological replicates. Scale bar =10 µm

**Figure 3:**
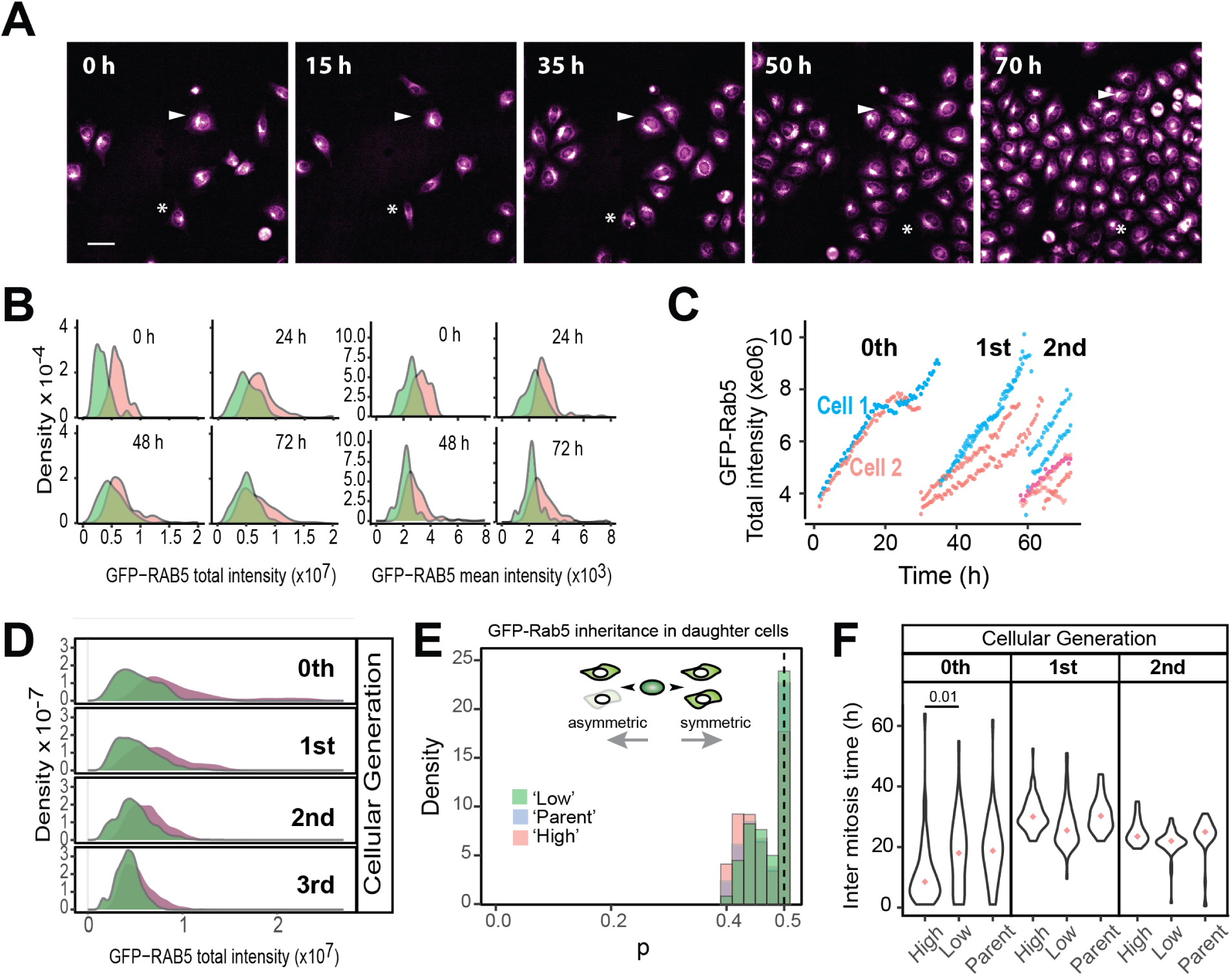
Time-lapse imaging reveals association between endocytic capacity and the cell cycle. **A)** Representative frames from time-lapse imaging of HeLa GFP-Rab5 cells. Movies were recorded every 30 minutes for 72 hours. Scale bar = 10 µm. **B)** Distributions of (i) total GFP-Rab5 intensity and (ii) mean GFP-Rab5 intensity normalized to cell area were quantified from the initial populations of 51 ‘low’ and 39 ‘high’ subpopulation cells at the indicated timepoints. **C)** Trajectory of total GFP-Rab5 intensity for representative cells across multiple rounds of cell division. **D)** Distribution of total GFP-Rab5 intensity in ‘low’ and ‘high’ subpopulations across multiple cell generations. **E)** Probability of asymmetric division between two daughter cells in parent, ‘low’, and ‘high’ subpopulations.**F)** Violin plot showing the inter-mitotic time for cells in parent, ‘low’, and ‘high’ subpopulations across different generations. Dots indicate population medians. p-values were computed using Wilcoxon Rank Sum (statistical test). Results are representative of two biological replicates.

We next sought to characterize the variation in endocytic capacity across other cell types. A panel of cell lines was pulsed with fluorescently tagged transferrin or dextran, and cell-to-cell variability was quantified by calculating the coefficient of variation in cargo uptake across individual cells within each population (Figure2D,E). All tested cell lines displayed a high degree of variability, indicating substantial differences in the endocytic capacities of individual cells. Notably, the coefficient of variation for dextran, a fluid-phase cargo, was consistently higher than that for transferrin, a receptor-mediated cargo, underscoring the more regulated nature of receptor-mediated endocytosis (Mayor et al., 2007). Furthermore, the two uptake modes exhibited variable correlations across different cell types (FigureS2A), suggesting cell type–specific mechanisms governing cargo internalization. Despite these differences, the consistently high variability observed for both cargoes across diverse cell lines demonstrates that heterogeneity in endocytic capacity is a fundamental property of cell populations.

Similar to macrophages, HeLa cells displayed high cell-to-cell variability in endocytosis. To test whether early endosomal content and cargo uptake were similarly associated, as observed in RAW cells, we pulsed HeLa cells stably expressing GFP–Rab5C under its endogenous promoter and regulatory elements (Poser et al., 2008) with fluorescently tagged transferrin or dextran and measured the correlation between cargo uptake and GFP–Rab5 levels. A strong positive correlation was observed for both cargoes (FigureS2B). To further characterize endocytic capacity, HeLa GFP–Rab5 cells were sorted into ‘high’ and ‘low’ subpopulations based on the GFP–Rab5 expression. Immunostaining confirmed corresponding differences in the number of endosomes and the total cellular intensities of Rab5 and its effector EEA1 between the two groups (Figure2F). We next asked whether these ‘high’ and ‘low’ subpopulations were stable or dynamic, as previously seen in RAW macrophages. After three days in culture, the initially distinct subpopulations had converged, becoming indistinguishable from the parental distribution (Figure2G). To further confirm this dynamic behavior, single-cell clones were derived from the pool of HeLa BAC GFP–Rab5 cells. In five independent clones, the initially distinct ‘high’ and ‘low’ endocytic capacities again converged to the parental distribution within four days (FigureS2C). Collectively, these findings demonstrate that endocytic capacity is a dynamic property of individual cells and that such variation is broadly conserved across mammalian cell types.

### Single cell analysis for drivers of heterogeneity of endocytic capacity

To better resolve the dynamic nature of endocytic capacity at the single-cell level, we performed live-cell time-lapse imaging of HeLa GFP–Rab5 cells (Figure3A, Movie S1A) for 72 hours at 30-min intervals. A custom image analysis pipeline was developed to segment and track single cells across multiple generations, combining Fiji (Schindelin et al., 2012), CellPose (Stringer et al., 2021; Pachitariu et al., 2022), and Ilastik (Sommer et al., 2011) (FigureS3A, Movie S1B). Since total GFP-Rab5 intensity correlates with cargo uptake (Figure2E), we used it as a metric for assessing the endocytic capacity of individual cells. Intensity profiles of single cells showed a steady increase in GFP-Rab5 levels over the cell’s lifespan, followed by a reset upon cell division (Figure3C). To test this behaviour across the endocytic subpopulations, we sorted cells into ‘low’ and ‘high’ subpopulations based on the GFP-Rab5 expression levels and performed live-cell imaging.

We then examined the distribution of total GFP-Rab5 intensity and mean intensity (area normalized) at the start of the imaging period and at defined time intervals (Figure3B). Analysis of the distribution of these parameters over a large number of cells showed that these parameters were initially distinct between ‘high’ and ‘low’ subpopulations, but eventually converged over time as the cells progressed through the cell cycle. Similar results were obtained in an unsorted population, where ‘high’ and ‘low’ groups were defined from the first frame of imaging (FigureS3B), ruling out cell sorting–associated artifacts. These findings were further corroborated by tracking GFP–Rab5 intensity across cell lineages over multiple generations (Figure3D). Together, live-cell imaging results are consistent with flow cytometry snapshots (Figure2G), reinforcing that endocytic capacity is dynamic: initially distinct subpopulations converge toward the parental distribution over successive cellular generations.

We next investigated potential causes of the heterogeneity in endocytic capacity observed in these images. One possibility is asymmetric distribution of organelles during mitosis, which has been reported to contribute to cell-to-cell variability (J. T. Chang et al., 2007; A. Y. Chang et al., 2017; Loeffler, Wehling, et al., 2019; Loeffler, Schneiter, et al., 2022). To test whether inheritance of GFP–Rab5 differed between the ‘high’ and ‘low’ cells, we measured GFP–Rab5 intensity in daughter cells immediately after division and assessed the probability of symmetric inheritance in the two subpopulations. The results (Figure3D) showed that GFP–Rab5 inheritance was largely symmetric in both ‘high’ and ‘low’ cells, ruling out asymmetric partitioning of GFP–Rab5 as a major contributor to population-level heterogeneity.

We next examined whether differences in inter-mitotic times could account for variation in endocytic capacity. To this end, we analyzed the inter-mitotic times of the ‘high’ and ‘low’ endocytic subpopulations across successive generations (Figure3F). The first—but not the subsequent—inter-mitotic time was markedly shorter for ‘high’ cells compared to ‘low’ cells. In two independent experiments, division time decreased from 16.16 h to 8.61 h (4̃7% reduction) and from 18 h to 8.5 h (5̃3% reduction), respectively. These findings indicate that immediately after sorting, ‘high’ endocytic cells were enriched in a population closer to mitosis than ‘low’ cells. Thus, sorting based on GFP–Rab5 intensity appears to preferentially enrich for cells in the G2 phase of the cell cycle, consistent with elevated GFP–Rab5 levels observed in single-cell traces preceding mitosis. Taken together, these results suggest that cell-cycle progression underlies heterogeneity in endocytic capacity, generating variability between individual cells at a given time snapshot.

### Cell cycle progression is a major driver of heterogeneity of endocytic capacity

Although live-cell imaging suggested a progressive increase in GFP–Rab5 intensity during the cell cycle, the stability of GFP raises the possibility that this trend reflects cumulative accumulation rather than active regulation. To address this, we pulsed HeLa GFP–Rab5 cells with fluorescently labeled transferrin and measured both GFP–Rab5 expression and transferrin uptake in relation to DNA content by flow cytometry. Both parameters showed a significant positive correlation with DNA content (Figure 4A). Imaging assays similarly revealed correlations between DAPI staining and early endosomal content, assessed by GFP–Rab5 and EEA1 (Figure 4B); the lower correlation observed in imaging may reflect measurement in a single plane rather than the full cell volume. Classification of cells into G1, S, and G2 phases confirmed that G2-phase cells display significantly higher GFP–Rab5 levels and transferrin uptake compared to G1 and S-phase cells (Figure 4C). Together, these results indicate that the increase reflects a bona fide, cell cycle–dependent rise in endocytic capacity rather than an artifact of GFP stability.

**Figure 4:**
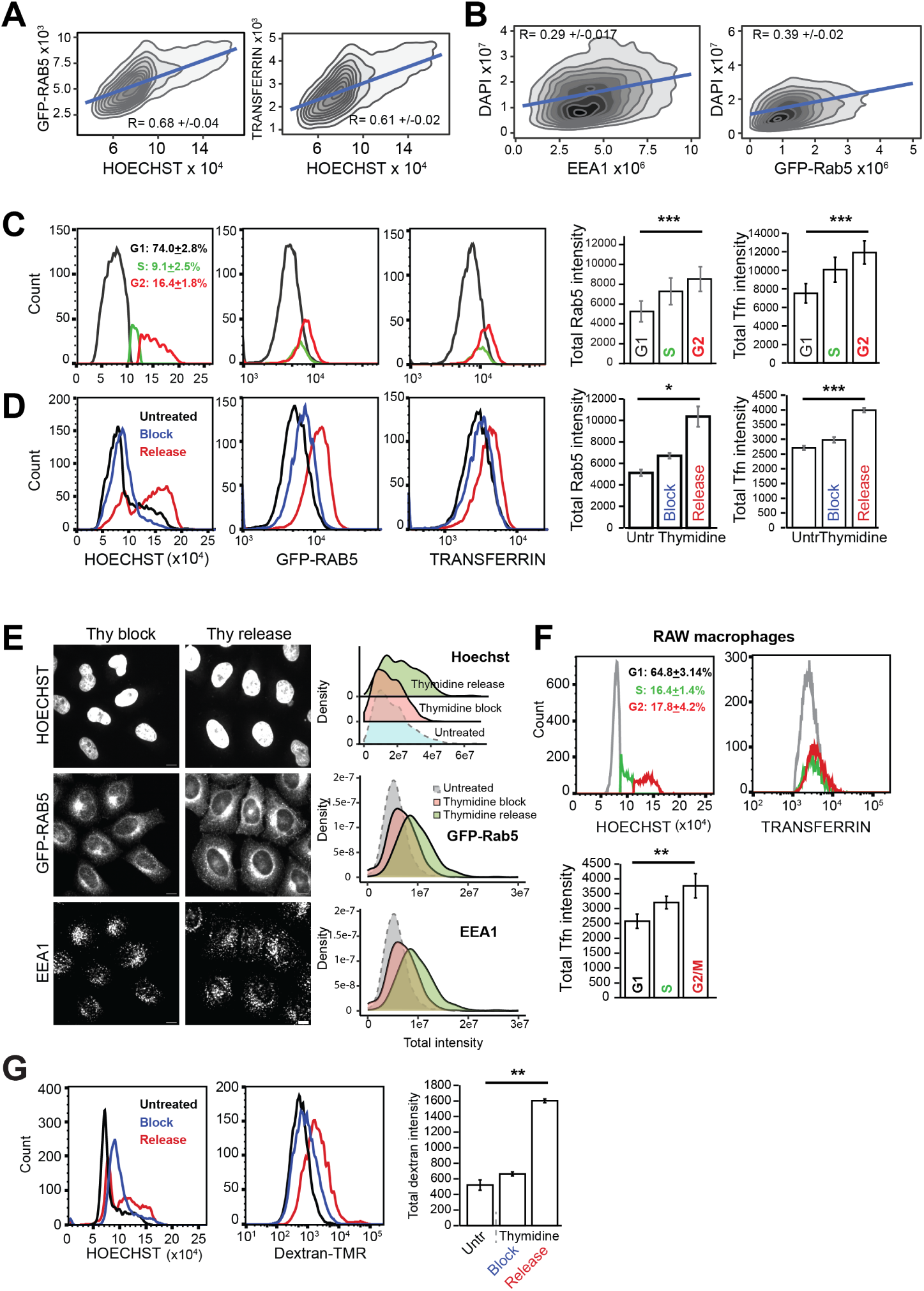
Cell cycle progression through interphase explains substantial variation in endocytic capacity. **A)** HeLa BAC GFP-Rab5 cells were pulsed with fluorescently labeled transferrin, followed by Hoechst staining and analyzed by flow cytometry. Contour plots show positive correlation between GFP-Rab5 (i) or transferrin (ii) and Hoechst intensity. **B)** HeLa GFP-Rab5 cells were fixed, immunostained for EEA1, and imaged. Contour density plots show relationships between DAPI intensity and total GFP-Rab5 (i) or EEA1 (ii) intensities. Data are representative of three independent experiments. For A, B, numbers denote mean ± SD of the correlation coefficients from three independent biological replicates.**C)** HeLa BAC GFP-Rab5 cells were pulsed with fluorescent transferrin, stained with Hoechst, and analyzed by flow cytometry. Cells were gated into G1, S, and G2/M phases based on Hoechst intensity. Bar graphs show mean total GFP-Rab5 and transferrin intensities within each gate. Error bars denote SD from four technical replicates. Data are representative of at least three independent biological replicates.**D, E)** HeLa BAC GFP-Rab5 cells were treated with 2.5 mM thymidine (Thy) for 18 hours to arrest the cell cycle in late G1/S phase. Cells at 0 hrs and 8 hrs post-treatment are labeled as block and release, respectively.**D)** Cells were pulsed with transferrin, fixed, and stained with Hoechst. Distributions of Hoechst, GFP-Rab5, and transferrin intensities are shown. Bar graphs display mean total GFP-Rab5 and transferrin intensities in each condition. Error bars indicate SD from two technical replicates. **E)**HeLa BAC GFP-Rab5 cells after thymidine block or release were fixed and immunostained for EEA1 to assess early endosomal content. Density plots from 450-1700 cells show distributions of nuclear, GFP-Rab5, and EEA1 intensities per cell. Data are representative of three independent experiments. (Scale bar = 10 µm).**F)** RAW 264.7 macrophages were pulsed with fluorescent transferrin, stained with Hoechst, and analyzed by flow cytometry. Cells were gated into G1, S, and G2/M phases. Bar graphs show mean transferrin intensity in each gate. Error bars represent SD from three technical replicates. **G)** RAW 264.7 macrophages were treated with 2 mM thymidine for 12 hours to arrest cells in late G1/S. Cells at 0 hrs and 4 hrs post-treatment (block and release) were pulsed with dextran, fixed, and stained with Hoechst. Distributions of Hoechst and dextran-TMR intensities are shown. Bar graphs depict mean total dextran-TMR intensity in each condition. Error bars represent SD from three technical replicates. For C-G, data are representative of at least three independent biological replicates.*denote *p*-values from Student’s *t*-test:\**p <* 0.05,\*\**p <* 0.01, \*\*\**p <* 0.001; ns, not significant.

To confirm these findings, HeLa GFP-Rab5 BAC cells were arrested at the G1/S boundary using thymidine treatment. Synchronous progression into G2 following release from the block enabled assessment of endocytic capacity in a G2-enriched cell population (Figure S4A). This experiment confirmed that an increased proportion of G2-phase cells leads to higher overall GFP–Rab5 intensity and transferrin uptake in the population (Figure 4D); similar results were obtained in two independent single-cell clones (Figure S4B,C). Further, thymidine-treated cells were immunostained for EEA1 and imaged, revealing a higher abundance of Rab5 and EEA1-positive endosomes in the G2-enriched condition following thymidine release (Figure 4E). Taken together, these results demonstrate that endocytic capacity fluctuates dynamically during interphase in HeLa cells, increasing as cells progress from G1/S to G2 phase.

We next investigated whether the relationship between cell cycle progression and endocytic capacity is conserved in macrophages by pulsing RAW macrophages with fluorescently labeled transferrin and assessing transferrin uptake in subpopulations stratified by DNA content. The results (Figure 4F) show that, similar to HeLa cells, RAW macrophages with higher DNA content exhibit increased endocytic capacity. To further validate this, cells were arrested with thymidine, briefly released from the block, and pulsed with fluorescently labeled dextran. This treatment enriched for G2-phase cells (Figure S4D), which displayed significantly higher dextran uptake (Figure 4G). Collectively, these findings demonstrate that as interphase cells progress from G1 to G2, endocytic uptake and early endosomal abundance increase continuously and reset after mitosis. Thus, cell cycle progression generates single-cell heterogeneity in endocytic capacity during interphase.

### Active control of endocytic capacity by interphase progression

To investigate sources of variation in endocytic capacity in macrophages using an independent approach, we performed single-cell RNA sequencing (scRNA-seq) of sorted ‘high’ and ‘low’ endocytic capacity RAW macrophage subpopulations (Figure 5A). The data were analyzed following current best practices for scRNA-seq analysis (Heumos et al., 2023) (Figure 5B). Gene ontology analysis of the top differentially expressed genes between the ‘high’ and ‘low’ subpopulations (Figure S5A) revealed significant enrichment of genes associated with cell division, mitosis, and G2-phase processes in the ‘high’ capacity cells. In contrast, the ‘low’ endocytic capacity subpopulation showed upregulation of genes related to transcriptional regulation, a process typically elevated in G1 phase (Bertoli et al., 2013) (Figure 5C,D). Comparison with cells following thymidine treatment, which enriches for G2-phase cells (Figure S5B), showed greater overlap of genes with the ‘high’ endocytic cells than with ‘low’ cells (Figure S5C). These findings provide additional evidence linking cell cycle progression to endocytic capacity. Further analysis of the scRNA-seq data revealed potential mechanisms beyond cell cycle contributing to endocytic heterogeneity. We overlaid differentially expressed genes (DEGs) between ‘high’ and ‘low’ endocytic subpopulations with DEGs from thymidine-treated versus parental populations (Figure 5Ei). As expected, cell cycle emerged as the most significantly enriched pathway among the intersecting genes (Figure 5Eii). We reasoned that the set of 205 non-overlapping genes could represent cell cycle–independent contributors to endocytic capacity. KEGG pathway analysis of this gene set showed significant enrichment in metabolic and lysosomal pathways. Together, these results establish cell cycle progression as a key determinant of endocytic capacity and identify additional cellular pathways contributing to its heterogeneity.

**Figure 5:**
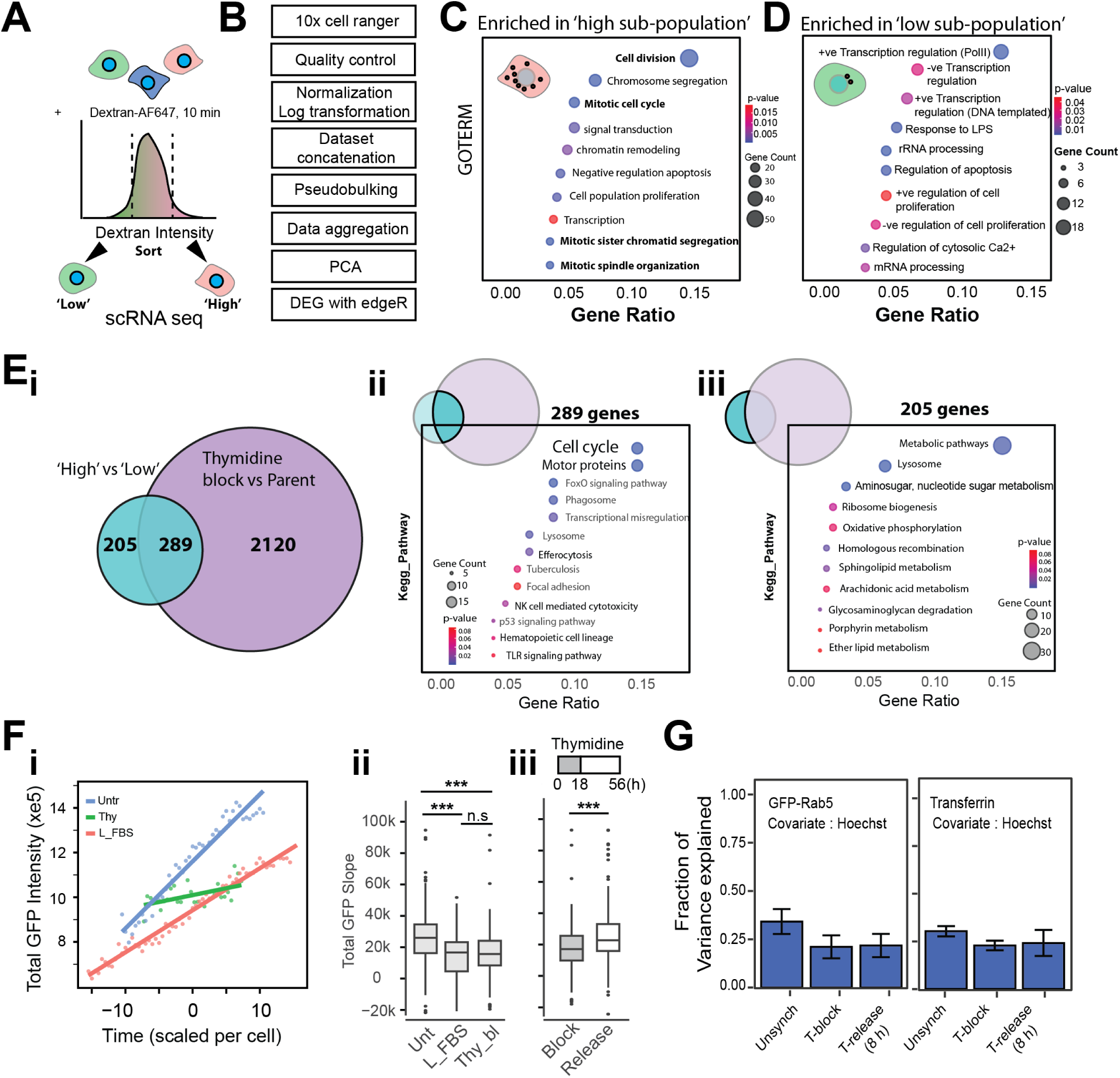
Cell cycle is a major contributing factor to endocytic heterogeneity in a population. **A)** Schematic of the single-cell RNA sequencing experiment. RAW 264.7 macrophages were pulsed with Dextran-TMR for 10 minutes and sorted into ‘low’ and ‘high’ endocytic subpopulations.**B)** Analysis pipeline used for scRNA-seq data.**C,D)** Pathway analysis from GO TERM enrichment showing biological processes enriched in genes upregulated in the ‘high’ (C) or ‘low’ (D) subpopulations. **E)** i) Venn diagram showing overlap of differentially expressed genes (DEGs) between ‘high’ vs ‘low’ endocytic capacity subpopulations, and DEGs from thymidine release (4 hrs) versus parent populations. ii) Pathway enrichment analysis of genes common to both. iii) Pathway enrichment of genes distinct to endocytic capacity.**F)** HeLa BAC GFP-Rab5 cells were imaged during thymidine (Thy) treatment or low FBS condition (1% FBS). Slopes were extracted from GFP-Rab5 intensity across time using a linear fit model from tracks with atleast 7 datapoints. i) Representative single cell tracks from the different conditions (dots) with the fits (solid lines). ii, iii) Comparison of slopes of total GFP-Rab5 cellular intensity between the indicated conditions. For panel iii, experimental conditions of thymidine block and release duration are shown in the figure. Slopes are calculated from 281, 56, 179, and 256 tracks for the experimental conditions denoted as Unt (untreated), L-FBS (low FBS), Thy bl (Thymidine Block) and release, respectively. *** and n.s. denote p-values *<* 0.0001, and *>* 0.1 respectively, from Student’s *t* -test.**G)** Variance decomposition analysis to quantify the contribution of cell cycle progression to endocytic heterogeneity. Error bars represent the standard deviation from the mean across three biological replicates.

While these results demonstrate a relationship between interphase progression and endocytic capacity, they do not exclude the possibility that endocytic capacity might increase constitutively in interphase cells, independent of active cell cycle–mediated regulation. To distinguish between these possibilities, we imaged HeLa GFP–Rab5 BAC cells under thymidine block. As an independent method to manipulate the cell cycle, cells were also grown in low FBS, a condition known to slow cell cycle progression (Orren et al., 1995; Qiuyan Chen et al., 2004). If endocytic capacity increases constitutively (i.e., independently of cell cycle progression), GFP–Rab5 intensity should continue to rise at a consistent rate despite cell cycle arrest. Conversely, if endocytic capacity is directly regulated by cell cycle progression, a decrease in the rate of GFP–Rab5 accumulation would be expected. To evaluate this, we fitted the GFP–Rab5 intensity from time-lapse measurements and calculated the slopes of these fits (Figure 5Fi). Comparison of slope distributions between non-synchronized and synchronized conditions (Figure 5Fii) revealed a significant reduction (∼30%) in the slope during both thymidine block and low FBS treatment. This indicates that cell cycle progression plays a direct role in modulating the heterogeneity in endocytic capacity. To further confirm this, we imaged the same cell population during and after release from thymidine block and assessed the slopes of the total GFP intensity. The results (Figure 5Fiii) showed that the initially low slope increased by ∼30% as cell cycle block was lifted and cells resumed growth. These findings confirm active control of total GFP–Rab5 content and endocytic capacity by cell cycle progression.

To quantify the precise contribution of cell cycle progression to the variation in endocytic capacity, we performed variance decomposition analysis on flow cytometry data (Figure 4D), as detailed in the Methods. The results (Figure 5G) show that approximately 30% of the variance in endocytic capacity - measured by either GFP-Rab5 content or transferrin uptake - can be explained by Hoechst staining, which serves as a marker of cell cycle progression. Thus, nearly one-third of the observed heterogeneity in endocytic capacity is attributable to interphase progression.

### Cell cycle progression promotes phenotypic diversity of *Mtb*

The identification of cell cycle progression as a key determinant of endocytic capacity suggests that it may likewise shape the phenotypic states of intracellular *Mtb*. To test this, we examined whether the phenotypic states of *Mtb* Mrx1-roGFP2 vary with the cell cycle phases of the infected host cell (Figure 6A). Asynchronous RAW cell populations were categorized into G1/S or G2 phases based on their DNA content, with gating performed using Draq-5 staining (Figure S6). Consistent with our earlier observations that G2-phase cells display higher endocytic capacity, and with prior report that cells with higher endocytic activity are more permissive to *Mtb* infection (Sachdeva et al., 2020), G2-phase cells exhibited higher infection rates than G1/S-phase cells (Figure 6Bi). Notably, a greater proportion of *Mtb* in G2-phase cells was oxidized, while the fraction in the reduced state decreased (Figure 6Bi). Together, these findings indicate that host cell cycle progression, by modulating endocytic capacity, influences the phenotypic state of intracellular *Mtb*.

**Figure 6:**
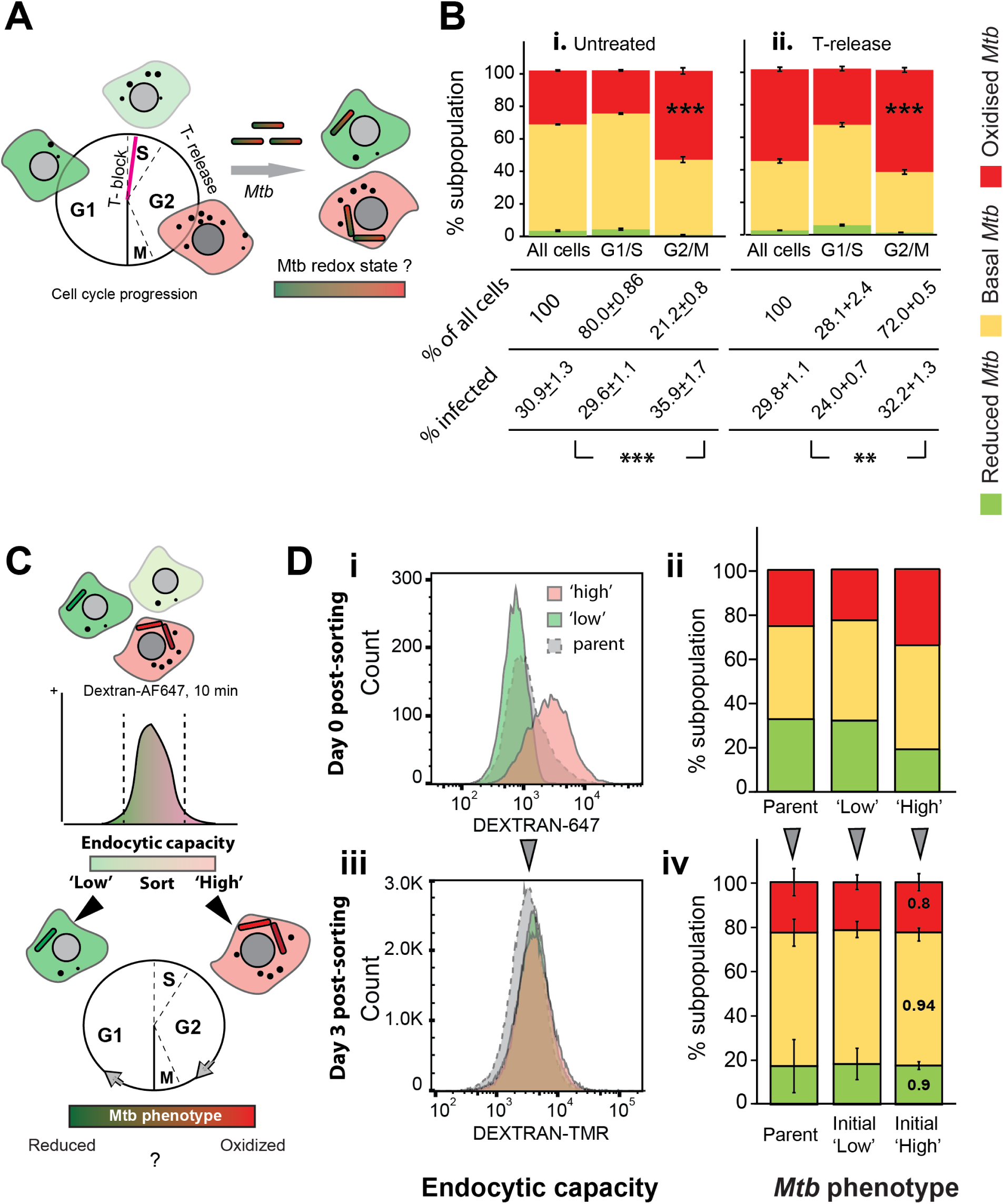
Cell cycle progression promotes phenotypic diversity of *Mtb*. **A**) Experimental scheme to assess the effect of cell cycle progression on intracellular *Mtb* phenotypic states. **B**) RAW 264.7 macrophages were treated with 2mM thymidine for 12 hrs for arresting the cell cycle in late G1/S phase. The cells were infected with *Mtb* Mrx1-roGFP2 H37Rv at 3.5 hrs post release, fixed and stained with Draq5. Graph shows the proportion of infected cells containing E_MSH_-reduced, basal and oxidised *Mtb* subpopulations between untreated or synchronised conditions (T-release) as indicated. Numbers denote the percentages of total and infected cells in each host cell subpopulations. **C**) Experimental scheme to assess plasticity of *Mtb* phenotypic states as cells move through the cell cycle. RAW 264.7 cells were infected with *Mtb* Mrx1-roGFP2 and pulsed with dextran for 10 mins in serum free media and sorted into ‘low’ or ‘high’ subpopulations based on the dextran fluorescence. **D**) Intracellular *Mtb* E_MSH_ subpopulations were analysed from 10,000 infected cells in the ‘low’ and ‘high’ endocytic subpopulations either immediately (i), or 2 days (ii) after sorting. Arrows indicate that the respective sorted subpopulations are cultured separately from each other. Error bars represents standard deviation of mean from 3 technical replicates, numbers denote *p*-value. Results are representative of atleast two independent biological replicates)(*denote *p*-values from Student’s *t*-test:\*\**p <* 0.01, \*\*\**p <* 0.001.)

To further validate this, cells were synchronized using thymidine block followed by release to enrich for G2-phase cells, infected with *Mtb* Mrx1-roGFP2, and the proportions of cells harboring oxidized, reduced, or basal *Mtb* were quantified. The results (Figure 6Bii) showed an increased proportion of cells with oxidized *Mtb* following thymidine treatment and in the G2-enriched subpopulation. Together, these data establish that cell cycle–dependent changes in endocytic capacity promote phenotypic diversity of intracellular *Mtb*.

Our previous result (Figure 3C) showed that endocytic capacity is reset as cells progress through mitosis (M phase). We therefore asked whether *Mtb* in ‘high’ endocytic capacity cells—experiencing higher oxidative stress—exhibits phenotypic plasticity as the host cell’s endocytic capacity changes following cell division. To test whether *Mtb* phenotypes adapt as host endocytic capacity changes, infected cells were sorted into ‘high’ and ‘low’ endocytic subpopulations, cultured separately for three days, and bacterial redox states were measured (Figure 6C). Once the initially distinct endocytic distributions had merged (Figure 6Di, iii), the proportion of oxidized *Mtb* in the former ‘high’ subpopulation decreased to parental levels, and no significant differences remained between the ‘high’ and ‘low’ groups (Figure 6Dii, iv). These results demonstrate that *Mtb* phenotypes exhibit plasticity in response to dynamic changes in host endocytic capacity as they transition from the G2 phase of the mother cell to the G1 phase of the daughter cells.

### Cell cycle driven changes in endocytic capacity and *Mtb* phenotypic diversity in post-differentiated macrophages

We next investigated whether the host cell cycle and endocytic capacity similarly shape *Mtb* phenotypes in terminally differentiated macrophages. First, we examined whether the relationship between endocytic capacity and cell cycle stage observed in RAW cells is conserved in THP-1 monocytes and the macrophages derived from them (schematic, Figure7A). THP-1 monocytes were pulsed with fluorescently labeled dextran and stained with Hoechst to distinguish cells in G1/S and G2 phases of the cell cycle. Endocytic capacity was quantified within these defined subpopulations. The same monocyte population was differentiated into macrophages using PMA stimulation, followed by analysis of both endocytic capacity and cell cycle distribution. The results (Figure7B) show that the proportions of G1/S and G2 cells remain comparable before and after differentiation, indicating that the cell cycle distribution of the monocyte population is largely maintained in the derived macrophages, consistent with previous reports (Gazova et al., 2020). The slight increase in G2-phase cells within macrophages may result from S-phase cells entering G2 during differentiation. Importantly, the relative differences in endocytic capacity between G1/S and G2 subpopulations persist following differentiation (Figure7B), demonstrating that cell cycle stages continue to influence endocytic capacity in differentiated cells.

Next, we assessed whether the phenotypic states of *Mtb* in differentiated macrophages exhibit similar dependence on endocytic capacity and cell cycle stage as observed in cycling cells. Macrophages (THP-1 or BMDM) infected with *Mtb* Mrx1-roGFP2 redox reporter were pulsed with fluorescently tagged dextran to delineate host subpopulations with ‘high’ or ‘low’ endocytic capacities, and stained with Draq5 to determine DNA content and cell cycle stage (FigureS7). The distribution of *Mtb* redox phenotypes was then assessed across these host subpopulations. The results (Figure7C–F) reveal that, similar to cycling cells, macrophages with higher endocytic capacity exhibit greater *Mtb* infection rates and harbor a higher proportion of oxidized *Mtb* compared to subpopulations with lower endocytic capacity (Figure 7C, E). Likewise, cells in the G2 phase are more infective and contain a greater proportion of oxidized *Mtb* than those in G1 (Figure7D, F). We further stratified the data based on oxidized or reduced *Mtb* subpopulations and analyzed the distribution of host macrophages across cell cycle stages and endocytic capacities. The results (Figure7G–J) show that oxidized *Mtb* predominantly associate with macrophage subpopulations exhibiting higher endocytic capacity, whereas reduced *Mtb* are more frequently found in cells with lower endocytic activity (Figure7G, I) in THP-1 monocyte derived macrophages as well as bone marrow derived macrophages. Similarly, reduced *Mtb* are enriched in G1-phase macrophages, while G2-phase macrophages predominantly harbor oxidized *Mtb* (Figure7H, J). These findings demonstrate that despite the absence of active cycling, terminally differentiated macrophages retain cell cycle stage–associated differences in endocytic capacity and influence corresponding *Mtb* phenotypic states.

**Figure 7:**
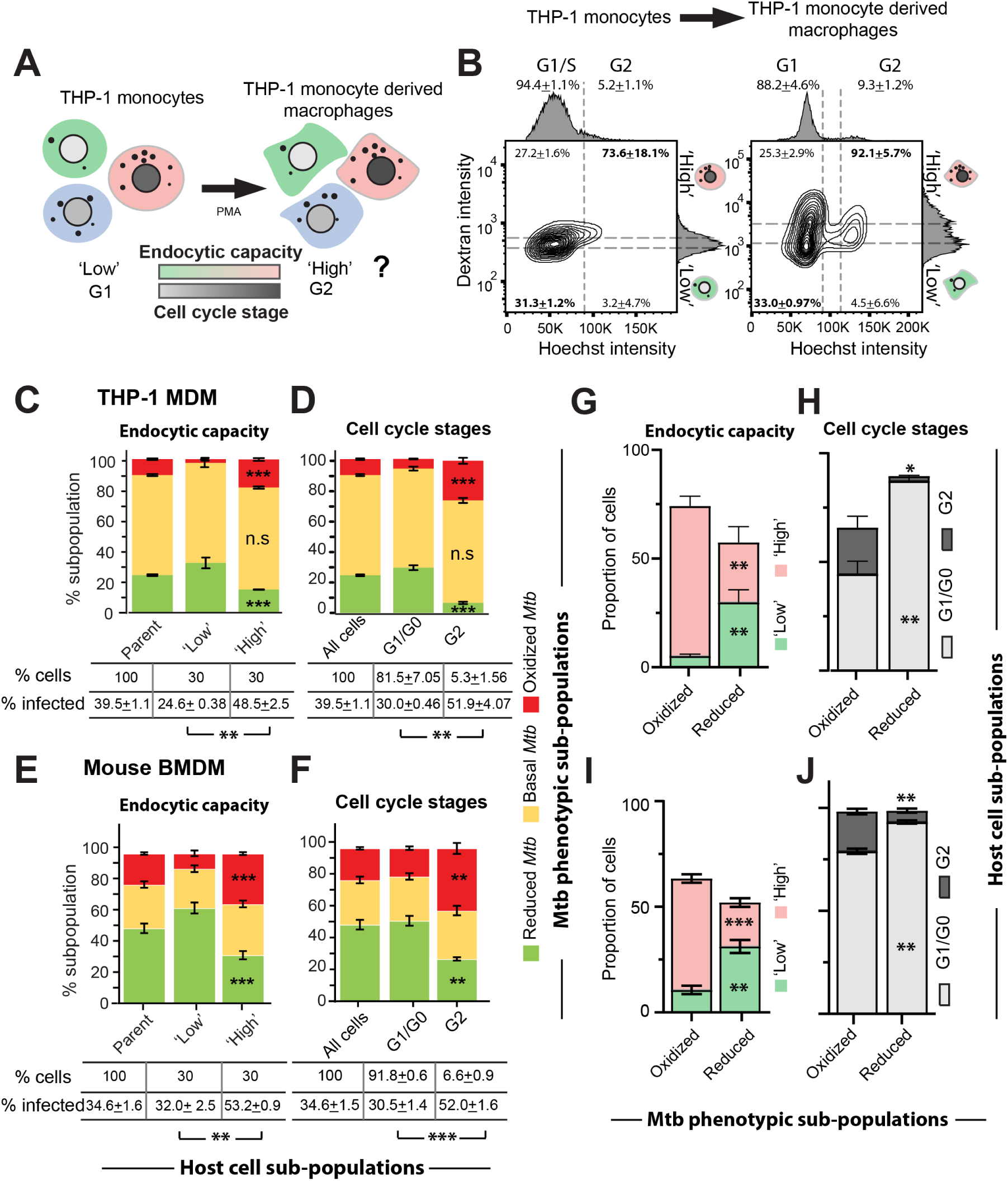
Interaction between bacterial and host heterogeneity in post differentiated macrophages. **A**) Schematic detailing the experiment to assess the relationship between endocytic capacity and cell cycle stages in macrophages post-differentiation. **B**) 2D FACS contour plot showing the endocytic capacity and cell cycle stages of THP-1 monocytes and the macrophages derived from them. To generate these plots, cells were pulsed with dextran for 10mins, fixed and stained with Hoechst 33342 (2 µg/mL for 20 min). Cells were gated into G1/S and G2/M subpopulations based on the Hoechst intensity. Proportions of cells in these gates are indicated. The subpopulations of the lower and upper 30% bounds of dextran intensity were defined as ‘low’ and ‘high’ dextran subpopulations, respectively. Numbers inside the plots indicate the proportion of G1/S and G2/M cells within the ‘low’ and ‘high’ dextran gates, and are expressed as mean ± standard deviation from three independent experiments. **C, D, E, F**) Proportion of different phenotypic subpopulations of *Mtb* Mrx1-roGFP2 in THP-1 macrophage (C,D) or BMDM (E,F) binned based on host cell endocytic capacity (C,E) or cell cycle stages (D,F). For these experiments, cells were infected with *Mtb* Mrx1-roGFP2 followed by pulsing with dextran for 10 mins, fixing, staining with DRAQ5 and measuring the endocytic capacity, cell cycle stages and redox of status of intracellular *Mtb* in the same cell population by flow cytometry. Numbers indicate the mean +/- SD of percentage infected cells from atleast three technical replicates. **G,H, I, J**) Among the E_MSH_-oxidized and reduced subpopulations, the percentage of bacteria in ‘low’ and ‘high’ dextran populations from THP-macrophages (G) or bone marrow derived macrophages (I), or cells in G1/G0 and G2 stages from THP-1 (H) or bone marrow derived macrophages (J) was quantified and represented as bar graphs. All data represents mean ± standard deviation of atleast three technical replicates in each subpopulation, and are representative of 3 independent biological experiments.

## Discussion

Cellular heterogeneity is a hallmark of biological systems (Altschuler et al., 2010; Komin et al., 2017), yet the mechanisms generating this variability and its impact on host-pathogen interactions remain incompletely understood. Here, we define a causal axis linking intrinsic heterogeneity in a fundamental host cellular process to intracellular bacterial phenotypic diversity. Using live-cell imaging, single-cell analyses, and bacterial reporter assays, we show that temporal shifts in endocytosis across interphase stages create structured single-cell variability, generating distinct intracellular niches that drive intracellular *Mtb* phenotypic adaptation.

These findings build upon our previous observation that cell-to-cell variability in endocytic capacity influences *Mtb* infectivity and subcellular trafficking (Sachdeva et al., 2020). We now demonstrate that this variability also shapes the intracellular microenvironments encountered by *Mtb* and alters its phenotypic adaptation (Figure1). We systematically characterize the distribution and temporal dynamics of endocytic capacity across cellular subpopulations in uninfected cells and show that this is not a static trait but fluctuates over time within individual cells (Figure2). This temporal variability suggests the presence of a structured source of heterogeneity (Huang, 2009; Movasat et al., 2025). Through live-cell imaging and single-cell RNA sequencing, we identify cell cycle progression as a key intrinsic determinant of endocytic capacity (Figure3). Cells in early and late interphase, i.e., G1 and G2 stages, exhibit marked differences in their endocytic capacities, and this variance aligns with changes in *Mtb* uptake and subsequent phenotypic adaptation during these stages. Our data suggest that the host cell cycle acts as a regulatory axis modulating the functional state of the endocytic machinery (FigureS4 and 5), thereby creating temporally dynamic intracellular niches for invading pathogens. This link between interphase progression and endocytic control has striking consequences for the pathogen. We find that *Mtb* phenotypic diversity is significantly influenced by the host cell’s position in the cell cycle at the time of infection (Figure6). Notably, this phenomenon persists in post-differentiated macrophages, indicating that the underlying regulatory architecture is preserved beyond cycling precursor populations (Figure7). Our data suggest that cell-cycle history leaves a lasting imprint and continues to shape cellular behavior in mature innate immune populations even after differentiation. This view is consistent with recent evidence indicating that macrophage functional diversity is maintained through both proliferative history and microenvironmental cues, rather than being erased by terminal differentiation. (Cao et al., 2024; Ahmad et al., 2022; Lavin et al., 2014). Together, these findings position cell cycle progression as a key regulator of functional heterogeneity in macrophages and identify endocytic capacity as a key effector of this variability. This dynamic heterogeneity, in turn, creates a mosaic of intracellular environments that drive bacterial phenotypic diversity.

Our findings resonate with several converging themes in cell and infection biology. They provide mechanistic insight into how functional heterogeneity arises within a cell population. While transcriptional noise, asymmetric division, and epigenetic regulation have been proposed as contributors to cellular heterogeneity (Carter et al., 2021; Cheow et al., 2016; Linker et al., 2019; Q. Chen et al., 2018; Harper et al., 2011; Rodriguez et al., 2019), our live-cell imaging and synchronization experiments identify interphase progression as a strong driver of variation in endocytic capacity. This supports emerging views of the cell cycle as a primary organizer of cellular state and function (Movasat et al., 2025; McDavid et al., 2016; Gruenheit et al., 2018). Our scRNA-seq data raises an intriguing possibility to explain active regulation of endocytosis by cell cycle progression: as cells prepare for volumetric doubling, membrane biogenesis must scale accordingly to account for expansion of the plasma membrane (Raucher et al., 1999) as well as internal compartments such as the ER. We speculate that endocytosis may act as a homeostatic buffer to calibrate plasma membrane composition and tension during volumetric expansion in interphase, thereby linking lipid biosynthesis, membrane dynamics, and trafficking capacity to cell cycle progression. Supporting this model, lipid biosynthesis genes are specifically upregulated in cells with high endocytic capacity.

Bacterial heterogeneity is a known driver of infection outcomes (Sherry et al., 2024; Aldridge et al., 2012; Dewachter et al., 2019), yet the contribution of host cell variability remains underexplored. Here, we establish that host endocytic capacity, which fluctuates with the cell cycle, is a critical factor shaping the intracellular niche. This capacity peaks in G2-phase, rendering cells more infectible. Crucially, we find that *Mtb* residing within these G2 cells display a more oxidized redox state, indicating they experience significant intracellular stress. Our scRNA-seq analysis provides the molecular explanation for this: high-uptake cells are characterized by an enriched lysosomal gene signature. This host-mediated pressure forces bacterial adaptation, structuring the phenotypic heterogeneity essential for persistence (Goossens et al., 2020; Mishra et al., 2019; Mavi et al., 2020). Our data demonstrates a remarkable phenotypic plasticity in the bacteria; as host cells divide and their daughters enter G1, their endocytic capacity resets, and we observe a corresponding reversion in the *Mtb* redox state. This shows that the host cell cycle continuously and dynamically structures the bacterial phenotypic diversity.

These results suggest that host cellular processes generate distinct cellular subpopulations that promote bacterial phenotypic diversification—a potential bet-hedging strategy in chronic infections. Our previous work demonstrated that cells with higher endocytic capacity are more permissive to *Mtb* entry but also more efficient at delivering internalized bacteria to lysosomes (Sachdeva et al., 2020) - posing a potential conundrum for *Mtb*. The pathogen appears to face a trade-off between infecting cells that are easier to enter but potentially more hostile intracellularly, versus cells that are harder to access but offer a more permissive intracellular environment. The current findings reinforce this trade-off by confirming that *Mtb* residing in high endocytic capacity or G2 subpopulations experience higher oxidative stress. Interestingly, our identification of cell cycle progression as a key driver of endocytic heterogeneity, alongside prior evidence that *Mtb* lipids can arrest host cell cycle at the G1/S boundary (Cumming et al., 2017), offers a potential resolution. It is plausible that *Mtb* exploits host cell cycle dynamics to its advantage by gaining entry into subpopulations with high endocytic activity or in G2 phase, then modulating the host cell cycle to maintain a G1-like state more conducive to its survival. The partial and heterogeneous nature of this arrest could create a dynamic cellular landscape where subpopulations of bacteria experience different intracellular conditions, facilitating phenotypic diversification or temporal regulation of stress exposure. Longitudinal single-cell tracking of both host and bacterial states will be essential to test this model and clarify the temporal interplay between *Mtb* physiology, endocytic capacity, and cell cycle states. In the broader context of infection biology, this study underscores the importance of considering host cell state as an active participant in shaping infection trajectories and microbial fate decisions.

Several important questions arise from these findings. First, what are the molecular regulators that link cell cycle progression to changes in the endocytic machinery? Our data point to Rab5 and EEA1 abundance and vesicle dynamics as effectors, but the upstream regulatory nodes remain to be defined. Second, how generalizable is the principle that cell cycle–dependent endocytic variation drives bacterial phenotypic adaptation? It will be important to test whether other intracellular pathogens, such as *Salmonella* or *Listeria*, experience similar variation in intracellular fate depending on host cell cycle or trafficking capacity. Third, does tissue-level variation in the distribution of host cell states (e.g., cycling versus quiescent macrophages) create spatial patterns of bacterial adaptation in vivo? Fourth, dissecting whether therapeutic modulation of cell cycle or endocytic pathways can bias bacterial populations toward more drug-sensitive states presents an exciting translational opportunity.

A key implication of our work is that, even in genetically identical cell populations, temporal progression through the cell cycle generates predictable fluctuations in pathogen permissiveness. This offers a new lens to interpret single-cell data from infections: what appears as stochastic heterogeneity may in fact emerge from structured host programs. By connecting single-cell variation in a host trait to phenotypic diversity in an intracellular pathogen, we provide a conceptual framework for understanding infection as an interaction between two dynamic, heterogeneous cellular subpopulations. Individual bacteria within this framework are likely to exhibit substantial differences in their susceptibility to antibacterial drugs (Sherry et al., 2024; Dhar and J. D. McKinney, 2007; Dewachter et al., 2019; Cotten et al., 2023). Identifying and defining such hotspots of drug tolerance within this matrix could provide a means to specifically target recalcitrant bacterial subpopulations. Indeed, *Mtb* of different redox states are known to differ in drug susceptibility (Bhaskar et al., 2014). Our findings identify host endocytic heterogeneity as a tunable parameter that shapes bacterial phenotypic landscapes. Modulating this axis—either by synchronizing host states or targeting pathways coupling cell cycle and endocytosis—may provide a route to bias *Mtb* populations away from stress-tolerant phenotypes and enhance drug efficacy.

In conclusion, we establish a direct mechanistic link between host cell cycle progression and the phenotypic landscape of an intracellular pathogen. By demonstrating that predictable host rhythms can structure bacterial diversity, this work offers a new lens for interpreting heterogeneity in hostpathogen interactions and provides a foundation for mapping these complex dynamics in vivo.

## Acknowledgments

We thank Amit Singh for the kind gift of the Mrx1-roGFP2 construct and, together with Dimple Notani, for critical reading of the manuscript. HeLa BAC GFP-Rab5 cells were kindly provided by Marino Zerial (MPI-CBG, Dresden). We also thank Vikas Yadav for assistance with *Mtb* redox data analysis and Prasanth Pillai for technical support. The following reagent was obtained through BEI Resources, NIAID, NIH: Macrophage Cell Line Derived from Wild Type Mice, NR-9456. We acknowledge the biosafety, central imaging and flow cytometry (CIFF), screening facilities, and the animal care and resource center (ACRC) at NCBS. The study was approved by the Institutional Animal Ethics and Biosafety Committees of NCBS. V.S. acknowledges funding from ANRF (CRG/2022/007379) and core support from NCBS-TIFR; N.S. acknowledges a fellowship from CSIR.

## Supplementary Information

### Supplementary Figures

**Figure S1:**
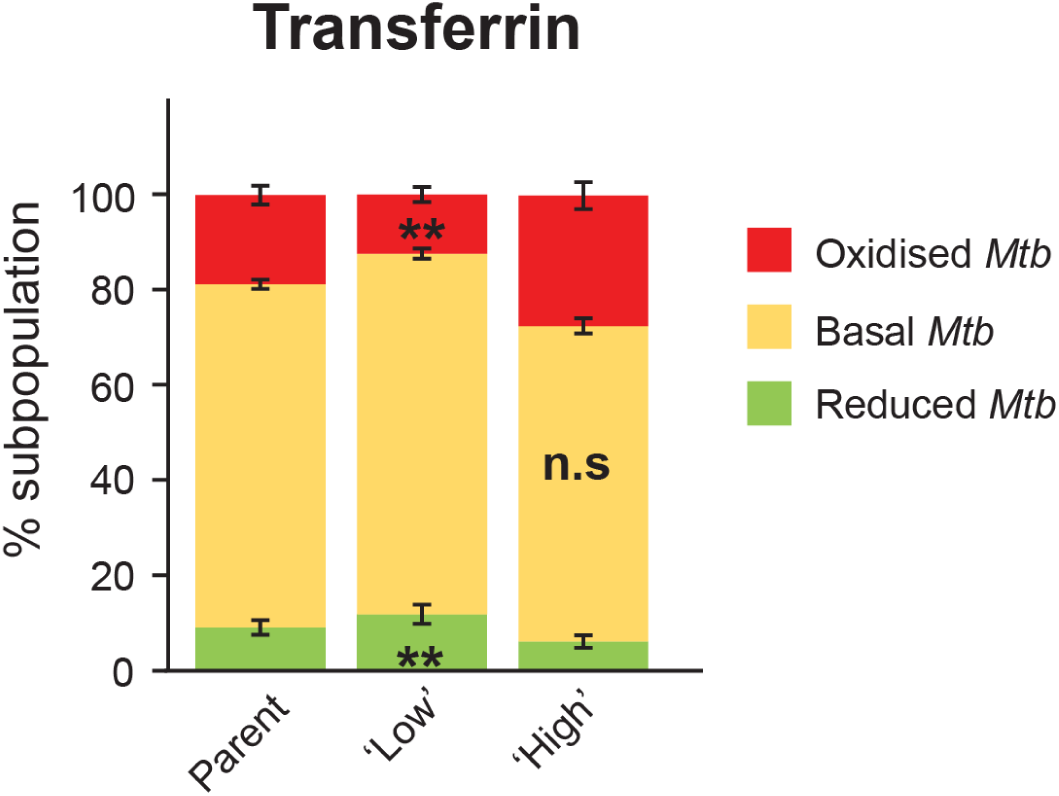
Heterogeneity in endocytic capacity promotes *Mtb* phenotypic diversity. (**A**) RAW 264.7 macrophages were infected with *Mtb* Mrx1-roGFP2 H37Rv for 2 hours at an MOI of 10 and pulsed with transferrin-AF647 for 10 minutes. Cells were fixed and analyzed by flow cytometry. Based on transferrin fluorescence intensity, infected cells were categorized into ‘high’ and ‘low’ endocytic capacity subpopulations. The redox status of intracellular *Mtb* was then assessed within these subpopulations. 10,000 infected cells were analyzed, and the top and bottom 30% of the transferrin fluorescence distribution were gated as ‘high’ and ‘low’ endocytic cells, respectively. The proportion of bacterial redox states (E_MSH_ reduced, basal, and oxidized) was determined as described in the Methods. The percentage of infected cells across these host subpopulations is indicated. Data shown are representative of at least three independent biological experiments. Error bars represent the standard deviation of technical replicates. *denote *p*-values from Student’s *t*-test:\*\**p <* 0.01, \*\*\**p <* 0.001; ns, not significant.

**Figure S2:**
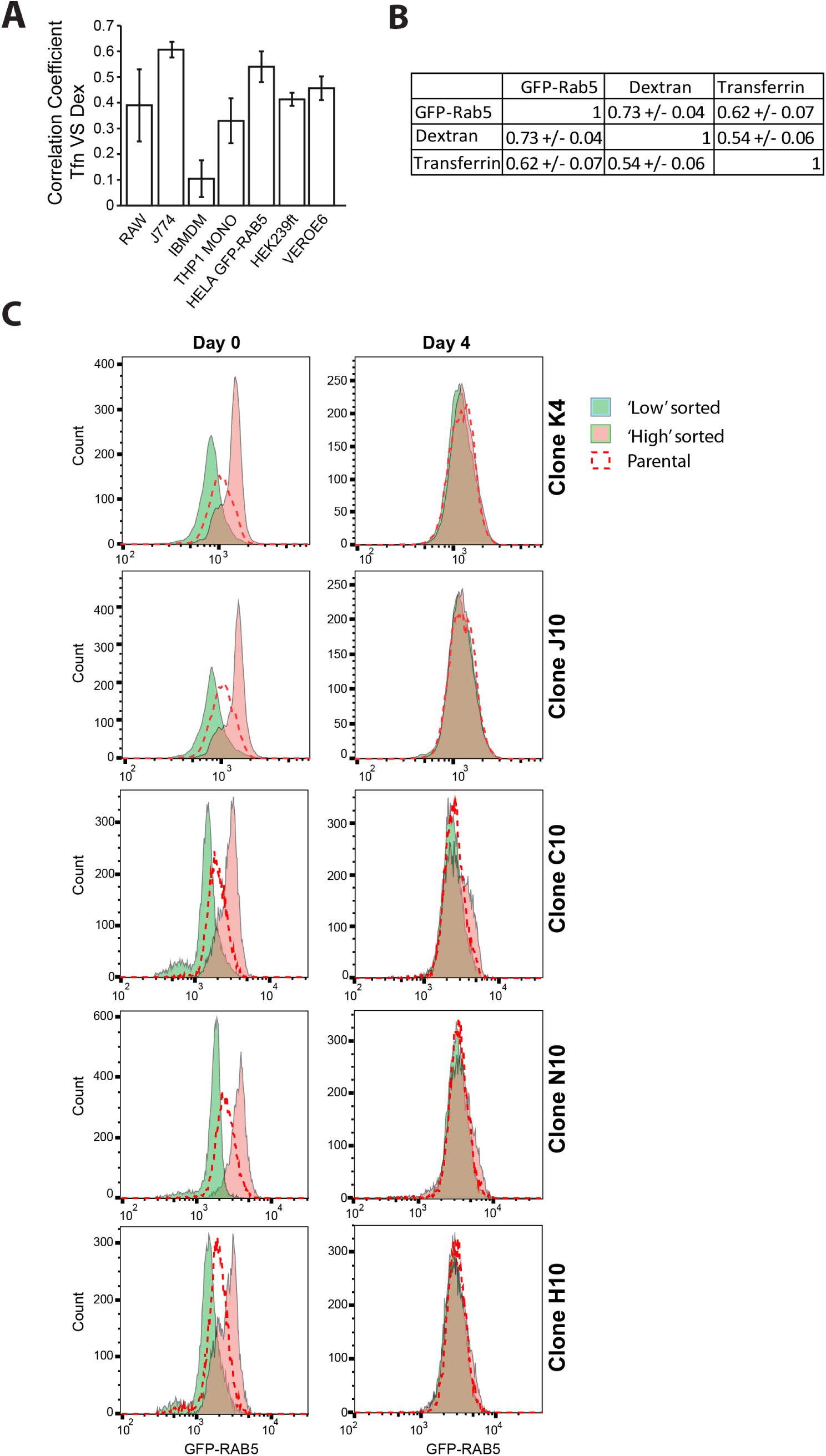
Characterization of variance in endocytic capacity across cell types. (**A**) Dextran and transferrin uptake assays were performed independently in the indicated cell lines. Correlation coefficients between dextran-TMR and transferrin-488 intensities were calculated from single-cell measurements. Error bars represent the standard deviation from the mean of three biological replicates.(**B**) Hela BAC GFP-Rab5 single-cell colonies were analysed after uptake of fluorescently tagged dextran and transferrin using flow cytometry. Heatmap shows the correlation coefficient between GFP-Rab5, dextran and transferrin. Data from three colnies are shown with error bars between colonies.(**C**) HeLa GFP-Rab5 single-cell colonies were sorted into ‘low’ and ‘high’ subpopulations based on GFP-Rab5 intensity. Shown are the distributions of GFP-Rab5 intensities from 10,000 cells in the ‘low’ and ‘high’ sorted HeLa GFP-Rab5 subpopulations either immediately after sorting (i, day 0) or four days post-sorting (ii, day 4). Red dots indicate the parental population. Distributions from five independent colonies are shown.

**Figure S3:**
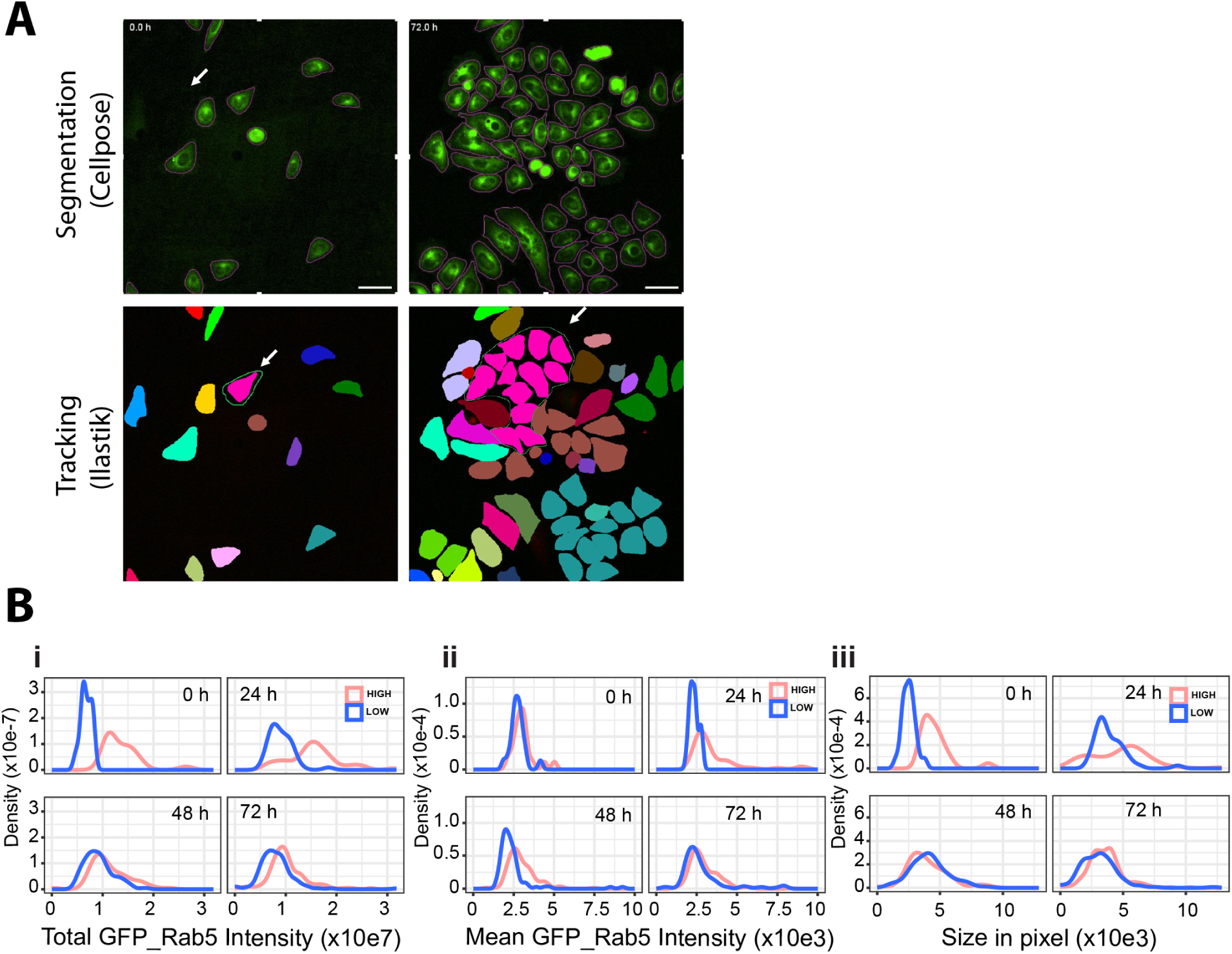
Time-lapse imaging reveals an association between endocytic capacity and the cell cycle. (**A**) Representative frames from time-lapse imaging of HeLa GFP-Rab5 cells used for segmentation and lineage tracking via Cellpose2 and ilastik, respectively. (**B**) Cells were binned into ‘high’ and ‘low’ groups (21 cells each) based on total GFP-Rab5 intensity of 84 cells from parental population at the start of imaging. Distribution analysis of these cell lineages across the indicated timepoints confirms that observed effects are not artifacts of the cell sorting procedure.

**Figure S4:**
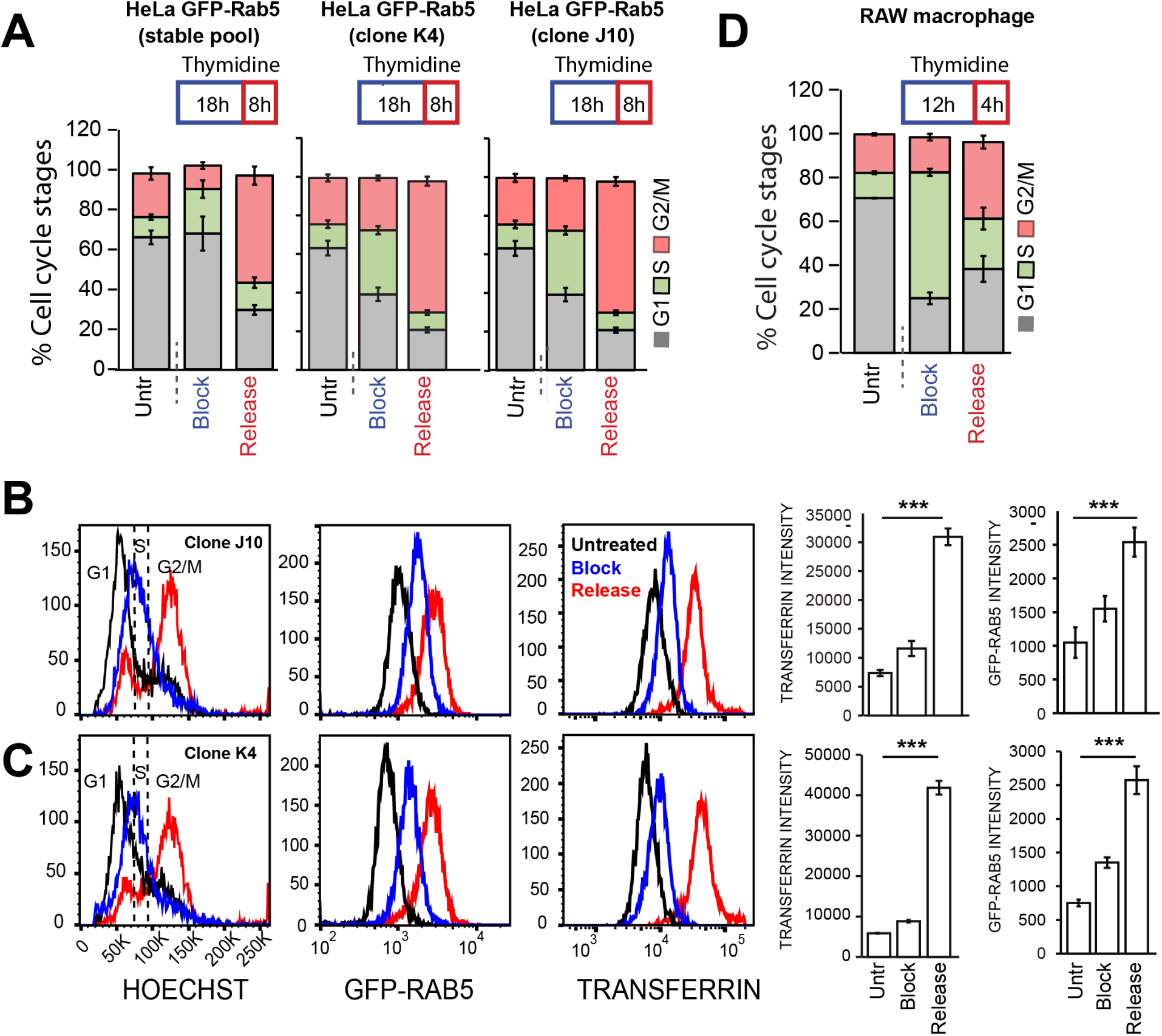
Cell cycle progression through interphase explains substantial variation in endocytic capacity. (**A**) HeLa BAC GFP-Rab5 cells were treated with 2.5 mM thymidine for 18 hours to arrest the cell cycle in late G1/S phase. Cells at 0 hours and 8 hours post-treatment are labeled as “block” and “release,” respectively. Cells were gated into G1, S, and G2/M phases based on Hoechst intensity. Bar graphs show the percentage of cells in each phase. Error bars denote standard deviation (SD) from four technical replicates. Data are representative of at least three independent biological replicates. (**B, C**) HeLa BAC GFP-Rab5 colonies K4 and J10 were subjected to a similar experiment. Cells were pulsed with transferrin, fixed, and stained with Hoechst. Distributions of Hoechst, GFP-Rab5, and transferrin intensities are shown. Bar graphs display the percentage of cells in different cell cycle stages. Error bars represent SD from three technical replicates. (**D**) RAW 264.7 macrophages were treated with 2 mM thymidine for 12 hours to arrest cells in late G1/S. Cells at 0 hours and 4 hours post-treatment (“block” and “release”) were pulsed with dextran, fixed, and stained with Hoechst. Bar graphs depict the percentage of cells in different cell cycle stages. Error bars represent SD from three technical replicates. Data are representative of at least three independent biological replicates. *denote *p*-values from Student’s *t*-test:\*\*\**p <* 0.001.

**Figure S5:**
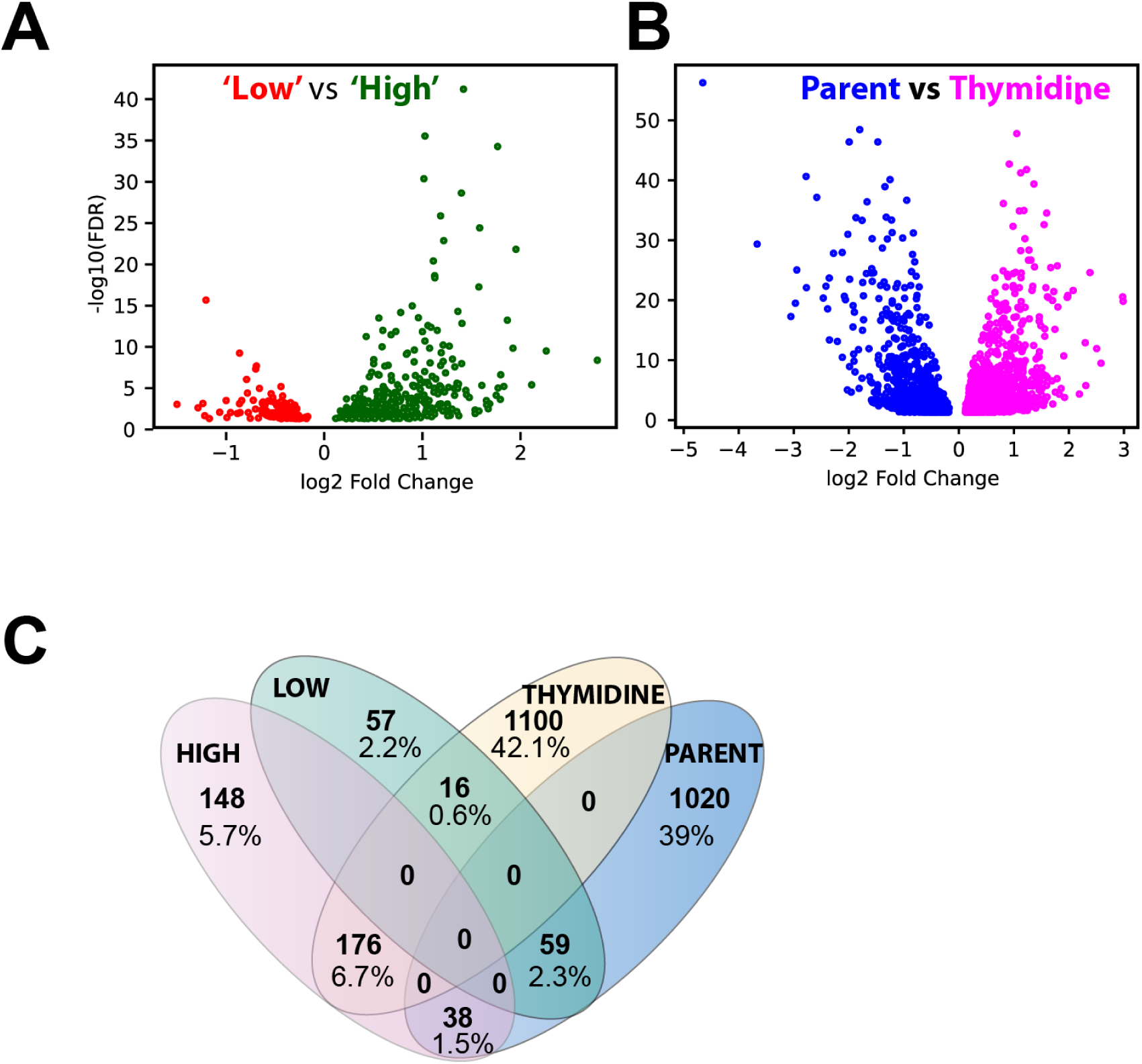
Cell cycle is a major contributing factor to endocytic heterogeneity. (**A,B**) Volcano plots showing differentially expressed genes between ‘low’ vs ‘high’ and parent vs thymidine release (4 h). (**C**) Venn diagram showing overlap of differentially expressed genes across conditions.

**Figure S6:**
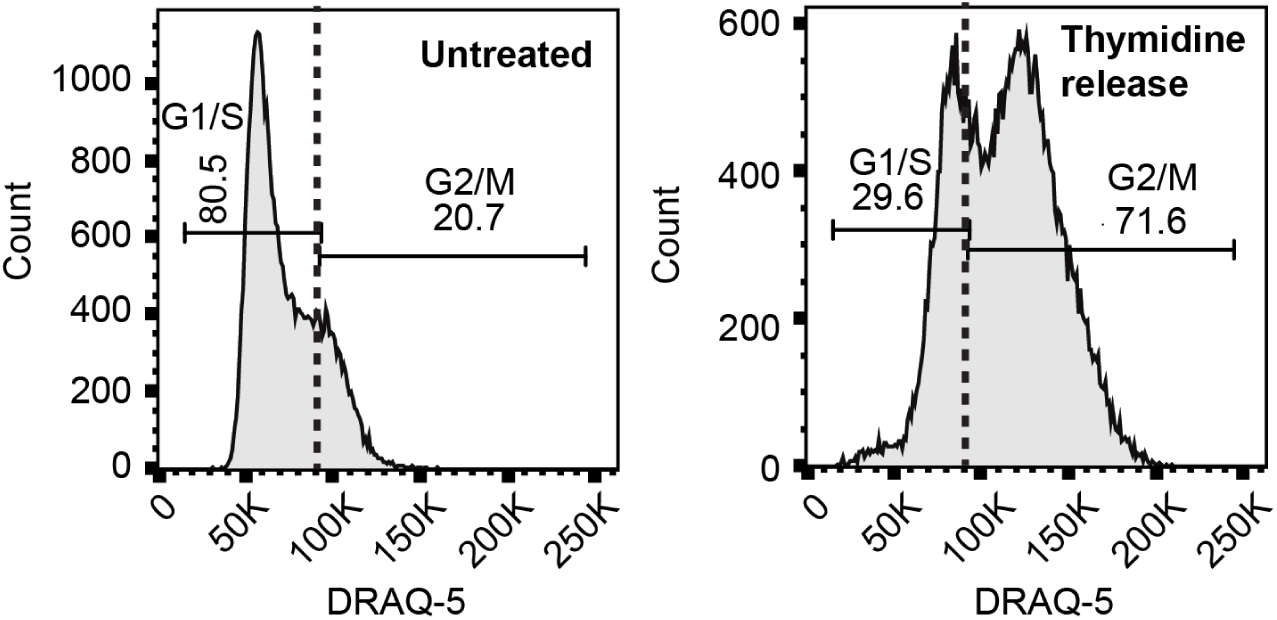
Cell cycle analysis using DRAQ5 in RAW cell synchronization. RAW 264.7 macrophages were treated with 2 mM thymidine for 12 h to arrest cells in late G1/S phase. Cells were then released into fresh medium for 3.5 hrs, followed by infection with *Mtb* Mrx1-roGFP2 H37Rv (MOI 20) for 30 min. After fixation and DRAQ5 staining, cell cycle stages were analyzed by flow cytometry. G1/S was gated based on the first sharp DRAQ5 intensity peak, while G2/M was gated at approximately twice the median intensity of G1.

**Figure S7:**
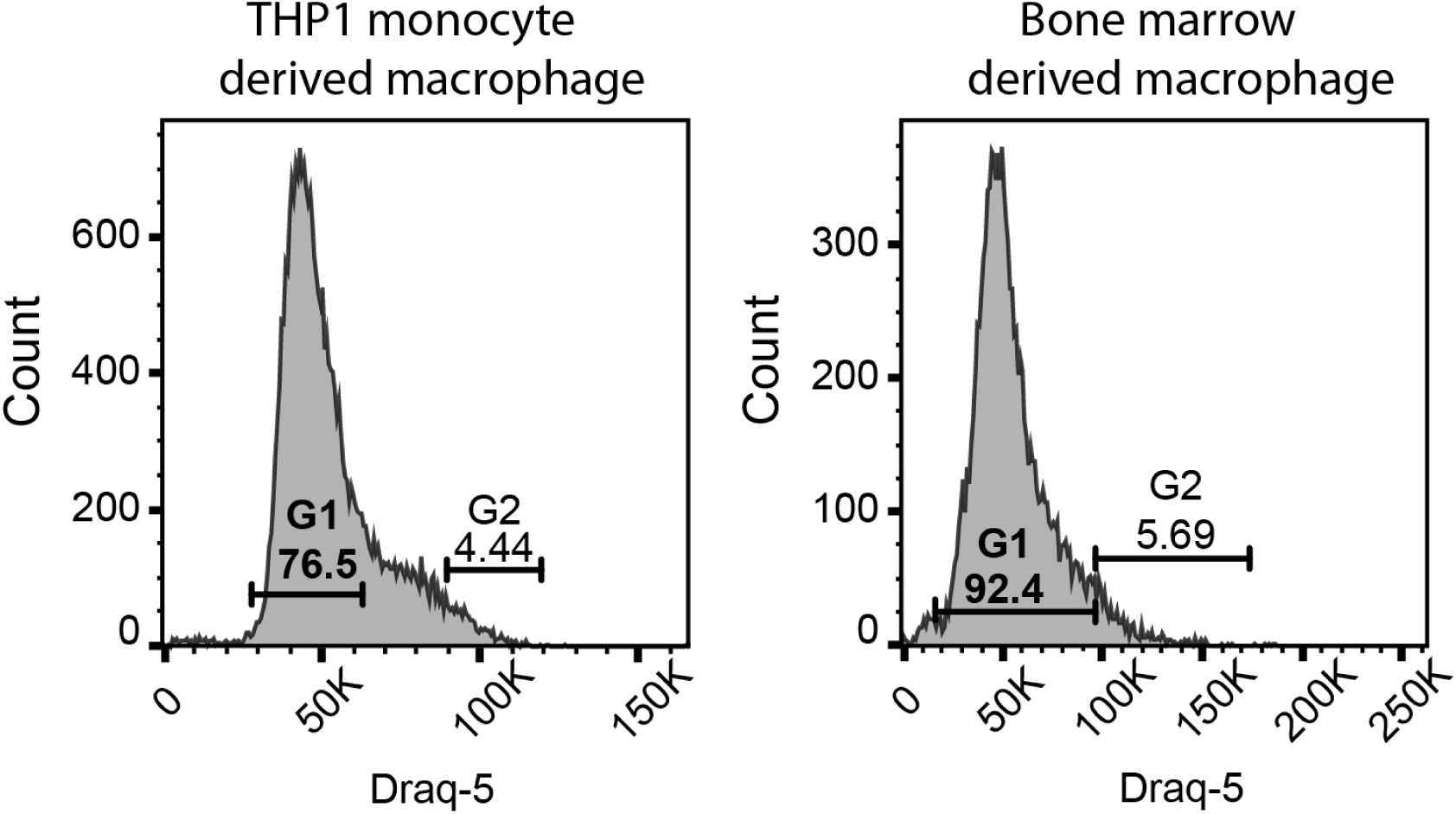
Cell cycle analysis using DRAQ5 in THP-1 and BMDM cells. THP-1 and BMDM cells were infected with *Mtb* Mrx1-roGFP2 at an MOI of 10, followed by a 10-minute dextran pulse, fixation, and staining with DRAQ5. Endocytic capacity and cell cycle stages were assessed by flow cytometry. Cell cycle phases were gated as indicated: the G1 gate was drawn based on the first sharp peak in DRAQ5 intensity; the G2 peak was gated using a threshold approximately twice the median intensity of G1.

## References

Adams, K. N., K. Takaki, L. E. Connolly, H. Wiedenhoft, K. Winglee, O. Humbert, P. H. Edelstein, C. L. Cosma, and L. Ramakrishnan (2011). “Drug tolerance in replicating mycobacteria mediated by a macrophage-induced efflux mechanism”. In: Cell 145.1, pp. 39–53.

Ahmad, Faraz, Anshu Rani, Anwar Alam, Sheeba Zarin, Saurabh Pandey, Hina Singh, Seyed Ehtesham Hasnain, and Nasreen Zafar Ehtesham (2022). “Macrophage: A Cell With Many Faces and Functions in Tuberculosis”. In: Frontiers in Immunology 13.5, pp. 1–18.

Akela, Ajit Kumar and Ashwani Kumar (June 2021). “Bioenergetic Heterogeneity in Mycobacterium tuberculosis Residing in Different Subcellular Niches.” In: mBio 12.3, e0108821.

Aldridge, Bree B, Marta Fernandez-Suarez, Danielle Heller, Vijay Ambravaneswaran, Daniel Irimia, Mehmet Toner, and Sarah M Fortune (Jan. 2012). “Asymmetry and aging of mycobacterial cells lead to variable growth and antibiotic susceptibility.” In: Science (New York, N.Y.) 335.6064, pp. 100–104.

Altschuler, Steven J. and Lani F. Wu (2010). “Cellular heterogeneity: Do differences make a difference?” In: Cell 141.4, pp. 559–563.

Avraham, R., N. Haseley, D. Brown, C. Penaranda, H. B. Jijon, J. J. Trombetta, R. Satija, A. K. Shalek, R. J. Xavier, A. Regev, and D. T. Hung (2015). “Pathogen cell-to-cell variability drives heterogeneity in host immune responses”. In: Cell 162.6, pp. 1309–1321.

Avraham, R. and D. T. Hung (2016). “A perspective on single cell behavior during infection”. In: Gut Microbes 7.6, pp. 518–525.

Baxter, E.W., A.E. Graham, N.A. Re, I.M. Carr, J.I. Robinson, S.L. Mackie, and A.W. Morgan (2020). “Standardized protocols for differentiation of THP-1 cells to macrophages with distinct M(IFN*γ*+LPS), M(IL-4) and M(IL-10) phenotypes”. In: Journal of Immunological Methods 478, p. 112721.

Bertoli, Cosetta, Jan M. Skotheim, and Robertus A. M. De Bruin (2013). “Control of cell cycle transcription during G1 and S phases”. In: Nature Reviews Molecular Cell Biology 14.8, pp. 518–528.

Bhaskar, Ashima, Manbeena Chawla, Mansi Mehta, Pankti Parikh, Pallavi Chandra, Devayani Bhave, Dhiraj Kumar, Kate S. Carroll, and Amit Singh (2014). “Reengineering Redox Sensitive GFP to Measure Mycothiol Redox Potential of Mycobacterium tuberculosis during Infection”. In: PLoS Pathogens 10.1, e1003902.

Bucci, Cecilia, Robert G. Parton, Ian H. Mather, Henk Stunnenberg, Kai Simons, Bernard Hoflack, and Marino Zerial (1992). “The small GTPase rab5 functions as a regulatory factor in the early endocytic pathway”. In: Cell 70.5, pp. 715–728.

Cadena, A. M., S. M. Fortune, and J. L. Flynn (2017). “Heterogeneity in tuberculosis”. In: Nat Rev Immunol 17.11, pp. 691–702.

Cao, Liren, Xiaoyan Meng, Zhiyuan Zhang, Zhonglong Liu, and Yue He (2024). “Macrophage heterogeneity and its interactions with stromal cells in tumour microenvironment”. In: Cell and Bioscience 14.1, pp. 1–22.

Carter, B. and K. Zhao (2021). “The epigenetic basis of cellular heterogeneity”. In: Nat Rev Genet 22.4, pp. 235–250.

Chang, Amy Y. and Wallace F. Marshall (2017). “Organelles - Understanding noise and heterogeneity in cell biology at an intermediate scale”. In: Journal of Cell Science 130.5, pp. 819–826.

Chang, John T., Vikram R. Palanivel, Ichiko Kinjyo, Felix Schambach, Andrew M. Intlekofer, and Arnob Banerjee (2007). “Asymmetric T lymphocyte division in the initiation of adaptive immune responses”. In: Science 315.5819, pp. 1687–1691.

Chen, Q., J. Shi, Y. Tao, and M. Zernicka-Goetz (2018). “Tracing the origin of heterogeneity and symmetry breaking in the early mammalian embryo”. In: Nat Commun 9.1, p. 1819.

Chen, Qiuyan and A Catharine Ross (2004). “Retinoic acid regulates cell cycle progression and cell differentiation in human monocytic THP-1 cells”. In: Experimental Cell Research 297.1, pp. 68–81.

Cheow, L. F., E. T. Courtois, Y. Tan, R. Viswanathan, Q. Xing, R. Z. Tan, D. S. Tan, P. Robson, Y. H. Loh, S. R. Quake, and W. F. Burkholder (2016). “Single-cell multimodal profiling reveals cellular epigenetic heterogeneity”. In: Nat Methods 13.10, pp. 833–836.

Collinet, Claudio, Martin Stoter, Charles R Bradshaw, Nikolay Samusik, Jochen C Rink, Denise Kenski, Bianca Habermann, Frank Buchholz, Robert Henschel, Matthias S Mueller, Wolfgang E Nagel, Eugenio Fava, Yannis Kalaidzidis, and Marino Zerial (Mar. 2010). “Systems survey of endocytosis by multiparametric image analysis.” In: Nature 464.7286, pp. 243–249.

Cotten, Katherine L and Kimberly Michele Davis (2023). “Bacterial heterogeneity and antibiotic persistence: bacterial mechanisms utilized in the host environment”. In: Microbiology and Molecular Biology Reviews 87.4, e00174–22.

Cumming, Bridgette M., Md Aejazur Rahman, Dirk A. Lamprecht, Kyle H. Rohde, Vikram Saini, John H. Adamson, David G. Russell, and Adrie J.C. Steyn (2017). “Mycobacterium tuberculosis arrests host cycle at the G1/S transition to establish long term infection (PLoS pathogens (2017) 13 5 (e1006389))”. In: PLoS pathogens 13.7, e1006490.

Delince, M. J., J. B. Bureau, A. T. Lopez-Jimenez, P. Cosson, T. Soldati, and J. D. McKinney (2016). “A microfluidic cell-trapping device for single-cell tracking of host-microbe interactions”. In: Lab Chip 16.17, pp. 3276–3285.

Dewachter, Liselot, Maarten Fauvart, and Jan Michiels (Oct. 2019). “Bacterial Heterogeneity and Antibiotic Survival: Understanding and Combatting Persistence and Heteroresistance”. In: Molecular Cell 76.2, pp. 255–267.

Dey, Gautam, Gagan D. Gupta, Balaji Ramalingam, Mugdha Sathe, Satyajit Mayor, and Mukund Thattai (2014). “Exploiting cell-to-cell variability to detect cellular perturbations”. In: PLoS ONE 9.3, e90540.

Dhar, Neeraj, John McKinney, and Giulia Manina (2016). “Phenotypic Heterogeneity in Mycobacterium tuberculosis”. In: Microbiology Spectrum 4.6, 10.1128/microbiolspec.tbtb2–0021–2016.

Dhar, Neeraj and John D McKinney (2007). “Microbial phenotypic heterogeneity and antibiotic tolerance”. In: Current Opinion in Microbiology 10.1, pp. 30–38.

Gazova, Iveta, Lucas Lefevre, Stephen J. Bush, Sara Clohisey, Erik Arner, Michiel de Hoon, Jessica Severin, Lucas van Duin, Robin Andersson, Andreas Lengeling, David A. Hume, and Kim M. Summers (2020). “The Transcriptional Network That Controls Growth Arrest and Macrophage Differentiation in the Human Myeloid Leukemia Cell Line THP-1”. In: Frontiers in Cell and Developmental Biology 8.July, pp. 1–21.

Gingold, Julian A., Ed S. Coakley, Jie Su, Dung Fang Lee, Zerlina Lau, Hongwei Zhou, Dan P. Felsenfeld, Christoph Schaniel, and Ihor R. Lemischka (2015). “Distribution Analyzer, a methodology for identifying and clustering outlier conditions from single-cell distributions, and its application to a Nanog reporter RNAi screen”. In: BMC Bioinformatics 16.1, pp. 1–20.

Goossens, Sander N, Samantha L Sampson, and Annelies Van Rie (2020). “Mechanisms of Drug-Induced Tolerance in Mycobacterium tuberculosis”. In: Clinical Microbiology Reviews 34.1. Article e00141-20.

Gough, Albert H., Ning Chen, Tong Ying Shun, Timothy R. Lezon, Robert C. Boltz, Celeste E. Reese, Jacob Wagner, Lawrence A. Vernetti, Jennifer R. Grandis, Adrian V. Lee, Andrew M. Stern, Mark E. Schurdak, and D. Lansing Taylor (2014). “Identifying and quantifying heterogeneity in high content analysis: Application of heterogeneity indices to drug discovery”. In: PLoS ONE 9.7, e102678.

Gruenheit, Nicole, Katie Parkinson, Christopher A. Brimson, Satoshi Kuwana, Edward J. John-son, Koki Nagayama, et al. (2018). “Cell Cycle Heterogeneity Can Generate Robust Cell Type Proportioning”. In: Developmental Cell 47.4, 494–508.e4.

Harper, C. V., B. Finkenstadt, D. J. Woodcock, S. Friedrichsen, S. Semprini, L. Ashall, D. G. Spiller, J. J. Mullins, D. A. Rand, J. R. Davis, and M. R. White (2011). “Dynamic analysis of stochastic transcription cycles”. In: PLoS Biol 9.4, e1000607.

Helaine, S., J. A. Thompson, K. G. Watson, M. Liu, C. Boyle, and D. W. Holden (2010). “Dynamics of intracellular bacterial replication at the single cell level”. In: Proc Natl Acad Sci U S A 107.8, pp. 3746–3751.

Helaine, Sophie, Angela M. Cheverton, Kathryn G. Watson, Laura M. Faure, Sophie A. Matthews, and David W. Holden (2014). “Internalization of Salmonella by macrophages induces formation of nonreplicating persisters”. In: Science 343.6167, pp. 204–208.

Helaine, Sophie, Brian P. Conlon, Kimberly M. Davis, and David G. Russell (2024). “Host stress drives tolerance and persistence: The bane of anti-microbial therapeutics”. In: Cell Host and Microbe 32.6, pp. 852–862.

Heumos, Lukas, Anna C. Schaar, Christopher Lance, Anastasia Litinetskaya, Felix Drost, Luke Zappia, Malte D. Lucken, Daniel C. Strobl, Juan Henao, Fabiola Curion, Hananeh Aliee, Meshal Ansari, Pau Badia-i-Mompel, Maren Buttner, Emma Dann, Daniel Dimitrov, Leander Dony, Amit Frishberg, Dongze He, Soroor Hediyeh-zadeh, Leon Hetzel, Ignacio L. Ibarra, Matthew G. Jones, Mohammad Lotfollahi, Laura D. Martens, Christian L. Muller, Mor Nitzan, Johannes Ostner, Giovanni Palla, Rob Patro, Zoe Piran, Ciro Ramirez-Suastegui, Julio Saez-Rodriguez, Hirak Sarkar, Benjamin Schubert, Lisa Sikkema, Avi Srivastava, Jovan Tanevski, Isaac Virshup, Philipp Weiler, Herbert B. Schiller, and Fabian J. Theis (2023). “Best practices for single-cell analysis across modalities”. In: Nature Reviews Genetics 24.8, pp. 550–572.

Huang, Sui (2009). “Non-genetic heterogeneity of cells in development: More than just noise”. In: Development 136.23, pp. 3853–3862.

Kamentsky, Lee, Thouis R. Jones, Adam Fraser, Mark-Anthony Bray, David J. Logan, Katherine L. Madden, Vebjorn Ljosa, Curtis Rueden, Kevin W. Eliceiri, and Anne E. Carpenter (Feb. 2011). “Improved structure, function and compatibility for CellProfiler: modular high-throughput image analysis software”. In: Bioinformatics 27.8, pp. 1179–1180.

Kim, Kang Ho and Joel M Sederstrom (July 2015). “Assaying Cell Cycle Status Using Flow Cytometry.” In: Current protocols in molecular biology 111, pp. 28.6.1–28.6.11.

Komin, Niko and Alexander Skupin (2017). “How to Address Cellular Heterogeneity by Distribution Biology”. In: Current Opinion in Systems Biology 3, pp. 154–160.

Krieger, Teresa and Benjamin D. Simons (2015). “Dynamic stem cell heterogeneity”. In: Development 142.8, pp. 1396–1406.

Lavin, Yonit, Deborah Winter, Ronnie Blecher-Gonen, Eyal David, Hadas Keren-Shaul, Miriam Merad, Steffen Jung, and Ido Amit (2014). “Tissue-resident macrophage enhancer landscapes are shaped by the local microenvironment”. In: Cell 159.6, pp. 1312–1326.

Linker, S. M., L. Urban, S. J. Clark, M. Chhatriwala, S. Amatya, D. J. McCarthy, I. Ebersberger, L. Vallier, W. Reik, O. Stegle, and M. J. Bonder (2019). “Combined single-cell profiling of expression and DNA methylation reveals splicing regulation and heterogeneity”. In: Genome Biol 20.1, p. 30.

Liu, Y., S. Tan, L. Huang, R. B. Abramovitch, K. H. Rohde, M. D. Zimmerman, C. Chen, V. Dartois, B. C. VanderVen, and D. G. Russell (2016). “Immune activation of the host cell induces drug tolerance in Mycobacterium tuberculosis both in vitro and in vivo”. In: J Exp Med 213.5, pp. 809–825.

Loeffler, Dirk, Florin Schneiter, Weijia Wang, Arne Wehling, Tobias Kull, Claudia Lengerke, Markus G. Manz, and Timm Schroeder (2022). “Asymmetric organelle inheritance predicts human blood stem cell fate”. In: Blood 139.13, pp. 2011–2023.

Loeffler, Dirk, Arne Wehling, Florin Schneiter, Yang Zhang, Niklas Müller-Bötticher, Philipp S. Hoppe, Oliver Hilsenbeck, Konstantinos D. Kokkaliaris, Max Endele, and Timm Schroeder (2019). “Asymmetric lysosome inheritance predicts activation of haematopoietic stem cells”. In: Nature 573.7774, pp. 426–429.

Mavi, Parminder Singh, Shweta Singh, and Ashwani Kumar (2020). “Reductive Stress: New Insights in Physiology and Drug Tolerance of Mycobacterium”. In: Antioxidants & Redox Signaling 32.18, pp. 1348–1366.

Mayor, Satyajit and Richard E Pagano (2007). “Pathways of clathrin-independent endocytosis”. In: Nature Reviews Molecular Cell Biology 8.8, pp. 603–612.

McDavid, Andrew, Greg Finak, and Raphael Gottardo (2016). “The Contribution of Cell Cycle to Heterogeneity in Single-Cell RNA-seq Data”. In: Nature Biotechnology 34.6, pp. 591–593.

McIntrye, J., D. Rowley, and G. R. Jenkin (1967). “The functional heterogeneity of macrophages at a single cell level”. In: Aust J Exp Biol Med Sci 45.6.

Mehta, Mansi, Raju S. Rajmani, and Amit Singh (2015). “Mycobacterium tuberculosis WhiB3 Responds to Vacuolar pH-induced Changes in Mycothiol Redox Potential to Modulate Phagosomal Maturation and Virulence”. In: Journal of Biological Chemistry 290.51, pp. 31005–31020.

Mills, E. and R. Avraham (2017). “Breaking the Population Barrier by Single Cell Analysis: One Host Against One Pathogen”. In: Current Opinion in Microbiology 36, pp. 69–75.

Mishra, Richa, Sakshi Kohli, Nitish Malhotra, Parijat Bandyopadhyay, Mansi Mehta, MohamedHusen Munshi, Vasista Adiga, Vijay Kamal Ahuja, Radha K Shandil, Raju S Rajmani, Aswin Sai Narain Seshasayee, and Amit Singh (2019). “Targeting redox heterogeneity to counteract drug tolerance in replicating Mycobacterium tuberculosis”. In: Science Translational Medicine 11.518, eaaw6635.

Monack, D. M., B. Raupach, A. E. Hromockyj, and S. Falkow (1996). “Salmonella typhimurium Invasion Induces Apoptosis in Infected Macrophages”. In: Proceedings of the National Academy of Sciences of the United States of America 93.18, pp. 9833–9838.

Mottet, M., C. Bosmani, N. Hanna, J. Nitschke, L. H. Lefrancois, and T. Soldati (2021). “Novel Single-Cell and High-Throughput Microscopy Techniques to Monitor Dictyostelium discoideum– Mycobacterium marinum Infection Dynamics”. In: Methods in Molecular Biology 2314, pp. 183–203.

Movasat, Hourieh, Enzo Giacopino, Ali Shahdoost, Yeganeh Dorri Nokoorani, Ali Houshyar Abrbekouh, Yaser Tahamtani, and Nika Shakiba (2025). “A Systems View of Cellular Heterogeneity: Unlocking the “Wheel of Fate””. In: Cell Systems, p. 101300.

Nagabhushanam, Vijaya, Alejandra Solache, Li-Min Ting, Claire J. Escaron, Jennifer Y. Zhang, and Joel D. Ernst (Nov. 2003). “Innate Inhibition of Adaptive Immunity: Mycobacterium tuberculosis-Induced IL-6 Inhibits Macrophage Responses to IFN-*γ*”. In: The Journal of Immunology 171.9, pp. 4750–4757.

Orren, David K, Lone N Petersen, and Vilhelm A Bohr (1995). “A UV-Responsive G2 Checkpoint in Rodent Cells”. In: Molecular and Cellular Biology 15.7, pp. 3722–3730.

Pachitariu, Marius and Carsen Stringer (2022). “Cellpose 2.0: How to Train Your Own Model”. In: Nature Methods 19.12, pp. 1634–1641.

Parbhoo, Trisha, Jacoba M. Mouton, and Samantha L. Sampson (2022). “Phenotypic Adaptation of Mycobacterium tuberculosis to Host-Associated Stressors That Induce Persister Formation”. In: Frontiers in Cellular and Infection Microbiology 12.

Peruzzi, Giovanna, Mattia Miotto, Roberta Maggio, Giancarlo Ruocco, and Giorgio Gosti (2021). “Asymmetric Binomial Statistics Explains Organelle Partitioning Variance in Cancer Cell Proliferation”. In: bioRxiv.

Poser, Ina, Mihail Sarov, James R.A. Hutchins, Jean Karim Hriche, Yusuke Toyoda, Andrei Pozniakovsky, Daniela Weigl, Anja Nitzsche, Bjorn Hegemann, Alexander W. Bird, Laurence Pelletier, Ralf Kittler, Sujun Hua, Ronald Naumann, Martina Augsburg, Martina M. Sykora, Helmut Hofe-meister, Youming Zhang, Kim Nasmyth, Kevin P. White, Steffen Dietzel, Karl Mechtler, Richard Durbin, A. Francis Stewart, Jan Michael Peters, Frank Buchholz, and Anthony A. Hyman (2008). “BAC TransgeneOmics: A high-throughput method for exploration of protein function in mammals”. In: Nature Methods 5.5, pp. 409–415.

Raucher, Drazen and Michael P. Sheetz (1999). “Membrane Expansion Increases Endocytosis Rate During Mitosis”. In: Journal of Cell Biology 144.3, pp. 497–506.

Rengarajan, Michelle and Julie A. Theriot (2020). “Rapidly Dynamic Host Cell Heterogeneity in Bacterial Adhesion Governs Susceptibility to Infection by Listeria monocytogenes”. In: Molecular Biology of the Cell 31.19, pp. 2097–2106.

Rodriguez, J., G. Ren, C. R. Day, K. Zhao, C. C. Chow, and D. R. Larson (2019). “Intrinsic Dynamics of a Human Gene Reveal the Basis of Expression Heterogeneity”. In: Cell 176.1–2, 213–226.e18.

Sachdeva, Kuldeep, Manisha Goel, and Varadharajan Sundaramurthy (2020). “Heterogeneity in the endocytic capacity of individual macrophage in a population determines its subsequent phagocytosis, infectivity and subcellular trafficking”. In: Traffic 21.8, pp. 522–533.

Saliba, Antoine-Emmanuel, Lei Li, Alexander J. Westermann, Silke Appenzeller, Daphne A. C. Stapels, Leon N. Schulte, Sophie Helaine, and Jorg Vogel (2016). “Single-Cell RNA-seq Ties Macrophage Polarization to Growth Rate of Intracellular Salmonella”. In: Nature Microbiology 2, p. 16206.

Schindelin, Johannes, Ignacio Arganda-Carreras, Erwin Frise, Verena Kaynig, Mark Longair, Tobias Pietzsch, Stephan Preibisch, Curtis Rueden, Stephan Saalfeld, Benjamin Schmid, Jean-Yves Tinevez, Daniel James White, Volker Hartenstein, Kevin Eliceiri, Pavel Tomancak, and Albert Cardona (July 2012). “Fiji: an Open-Source Platform for Biological-Image Analysis”. In: Nature Methods 9.7, pp. 676–682.

Sherry, Jessica and E. Hesper Rego (2024). “Phenotypic Heterogeneity in Pathogens”. In: Annual Review of Genetics 58.Volume 58, 2024, pp. 183–209.

Snijder, B., R. Sacher, P. Rämö, E. M. Damm, and P. Liberali (2009). “Population Context Determines Cell-to-Cell Variability in Endocytosis and Virus Infection”. In: Nature 461, pp. 520–523.

Sommer, Christoph, Christoph Straehle, Uwe Kothe, and Fred A. Hamprecht (2011). “Ilastik: Interactive Learning and Segmentation Toolkit”. In: 2011 IEEE International Symposium on Biomedical Imaging: From Nano to Macro, pp. 230–233.

Stringer, Carsen, Tim Wang, Michalis Michaelos, and Marius Pachitariu (2021). “Cellpose: A Generalist Algorithm for Cellular Segmentation”. In: Nature Methods 18.1, pp. 100–106.

Sukumar, N., S. Tan, B. B. Aldridge, and D. G. Russell (2014). “Exploitation of Mycobacterium tuberculosis Reporter Strains to Probe the Impact of Vaccination at Sites of Infection”. In: PLoS Pathogens 10.9, e1004394.

Tan, S., N. Sukumar, R. B. Abramovitch, T. Parish, and D. G. Russell (2013). “Mycobacterium tuberculosis Responds to Chloride and pH as Synergistic Cues to the Immune Status of Its Host Cell”. In: PLoS Pathogens 9.4, e1003282.

Tan, S., R. M. Yates, and D. G. Russell (2017). “Mycobacterium tuberculosis: Readouts of Bacterial Fitness and the Environment Within the Phagosome”. In: Methods in Molecular Biology 1519, pp. 333–347.

Thottacherry, Joseph Jose, Mugdha Sathe, Chaitra Prabhakara, and Satyajit Mayor (2019). “Spoilt for choice: Diverse endocytic pathways function at the cell surface”. In: Annual Review of Cell and Developmental Biology 35, pp. 55–84.

Toda, Gotaro, Toshimasa Yamauchi, Takashi Kadowaki, and Kohjiro Ueki (2021). “Preparation and culture of bone marrow-derived macrophages from mice for functional analysis”. In: STAR Protocols 2.1, p. 100246.

Zeigerer, Anja, Jerome Gilleron, Roman L. Bogorad, Giovanni Marsico, Hidenori Nonaka, Sarah Seifert, Hila Epstein-Barash, Satya Kuchimanchi, Chang Geng Peng, Vera M. Ruda, Perla Del Conte-Zerial, Jan G. Hengstler, Yannis Kalaidzidis, Victor Koteliansky, and Marino Zerial (2012). “Rab5 is necessary for the biogenesis of the endolysosomal system in vivo”. In: Nature 485.7399, pp. 465–470.

